# A fatty acid-binding protein links lipid handling to chitin synthase-dependent cuticle formation and nanoscale organization

**DOI:** 10.64898/2026.09.17.752344

**Authors:** Marius Beck, Kim Venus, Weixing Zhu, Luca Bertinetti, Yael Politi, Hans Merzendorfer

## Abstract

**Background:** The insect cuticle is a chitin-containing extracellular matrix whose formation requires precise coordination of chitin biosynthesis, fibril organization, and post-synthetic maturation. Although lipids are essential components of the cuticle, the molecular mechanisms linking lipid transport to chitin deposition remain poorly understood. Here we identify TcFabp8, a fatty acid-binding protein in *Tribolium castaneum*, as a functionally important regulator of chitin synthase 1 (Chs1)-dependent cuticle formation.

**Results:** RNAi-mediated depletion of *TcFABP8* caused molting defects, reduced cuticle thickness, loss of apical TcChs1 immunoreactivity, and reduced chitin deposition and organization. The finding that *TcCHS1* transcript levels were unaffected by RNAi targeting *TcFABP8* suggests that a post-transcriptional step, potentially involving vesicular TcChs1 transport, is affected, thereby impairing chitin synthesis and deposition. Synchrotron X-ray diffraction analysis of adult elytra revealed that TcFabp8 depletion preserved characteristic α-chitin-associated scattering features, but altered chitin fibril accumulation, crystalline coherence, and the spatially ordered alignment of fibrils within the cuticular matrix. The *TcFabp8* gene locus encodes splice variants yielding different N-terminal architectures: TcFabp8X1 carries a Sec/SPI signal peptide which is absent in TcFabp8X2 and TcFabp8X3. Immunocytochemistry using TcFabp8- specific antibodies detected intracellular signals in subapical cytoplasmic puncta and extracellular signals within the cuticle layers, which partially colocalized with Nile red fluorescence indicating lipid-rich environments.

**Conclusions:** Together, our data demonstrate that TcFabp8 is a critical facilitator of TcChs1-dependent chitin deposition and extends functional connection between lipid transport and chitin biosynthesis in insects.

## 1. Background

The insect cuticle is a chitin-containing extracellular matrix that provides mechanical protection, prevents desiccation, and enables locomotion. Its formation requires precise coordination between chitin polymer biosynthesis and post-synthetic maturation processes, including fibril organization, tanning, and sclerotization. In insects, chitin synthase 1 (Chs1) is essential for epidermal cuticle formation, while Chs2 is required for peritrophic matrix formation in the gut [1]. The distinct functions of insect *CHS* genes, first reported in *Manduca sexta* [2, 3], were ultimately confirmed through RNA interference experiments in *Tribolium castaneum* [4–6]. Moreover, these studies demonstrated the central role of *TcCHS1* in molting, survival, and adult cuticle integrity. In beetles such as *T. castaneum*, the forewings are evolutionarily reshaped to elytra, which are highly sclerotized cuticular structures that serve as armor plates and covering the delicate hindwings and dorsal abdomen. The exceptional rigidity and durability of the elytral cuticle are attributed to its organization into a multilamellar chitin-protein composite with extensive protein cross-linking resulting from quinone tanning [7].

After polymerization, single chitin chains are transported across the apical membrane through an extrusion pore of Chs1 finally reaching the assembly zone at the apical epidermal surface [8]. Within this compartment, nascent chitin chains assemble into nanofibrils that are bundled to thicker fibers, organized into flat sheets that are stacked on top of each other frequently in helicoidal arrangements to finally yield the multilayered ultrastructure of the cuticle. Genetic and ultrastructural analyses in *Drosophila melanogaster* identified Obstructor-A (ObstA) and Knickkopf (Knk) as key organizers that stabilize nascent chitin against premature degradation during this process [9–12]. Additional chitin-modifying enzymes including chitin deacetylases and cuticular chitinases, further remodel the polymer before it is organized into mechanically competent laminar architectures [13–15].

Beyond the 3D structural matrix organizers, several auxiliary factors have been shown to influence Chs1 localization and activity. In *D. melanogaster*, Chs1 physically associates with proteins such as Twinstar and SERCA involved in cytoskeletal dynamics and calcium homeostasis, supporting the view that actin organization, and Ca^2+^-regulated membrane trafficking contribute to efficient chitin deposition [16, 17]. These findings emphasize that Chs1-dependent cuticle formation is regulated by a multi-component process rather than by purely controlling enzyme activity [18]. Chitin synthesis, protein cross-linking, and other pathways, such as sclerotization which is hardening the cuticle by chemical crosslinking [7], are tightly regulated in space and time, as developing epidermal cells secrete the cuticle in a stratified fashion [19].

Next to chitin and proteins, the cuticle’s barrier function depends on lipid-derived components. The outermost epicuticular envelope is a waxy layer, rich in long-chained lipids. It waterproofs the cuticle and provides a barrier to xenobiotics. Disruption of cuticular lipid deposition can be lethal – for instance, RNAi mediated silencing of genes encoding the insect envelope protein “Snustorr snärlik” or the lipid-transporting ABC transporter ABCH-9C results in loss of surface lipids and fatal desiccation of larvae [20, 21]. Despite the clear importance of lipids in cuticle biology, the molecular mechanisms linking lipid transport and metabolism to cuticle formation remain poorly understood. Fatty acid-binding proteins (Fabps) are a family of small, evolutionarily conserved proteins that bind long-chain fatty acids and other hydrophobic ligands and function as intracellular lipid chaperones or transporters. In insects, Fabps have been reported from specialized tissues and are increasingly linked to developmental lipid handling and metabolic homeostasis. For example, juvenile hormone analogue exposure in *Locusta migratoria* strongly downregulates a midgut-enriched Fabp, and RNAi-mediated depletion of its gene reduces triglyceride levels and lipid droplet accumulation in the fat body [22–25]. In *Bombyx mori*, Fabp2/3 expressed in the midgut in a food-dependent manner were linked with fatty acid trafficking, metabolic routing, oxidative stress and larval body pattern pigmentation [26]. Recent work in *D. melanogaster* provided a first link between lipid transport and chitin synthesis in the integument by showing that Fabp physically associates with Chs1 and that Fabp depletion impairs wing chitin deposition and the cuticular barrier function [17]. Whether Fabp-dependent regulation of chitin biosynthesis is conserved in non-dipteran insects and whether it contributes to adult cuticle architecture remains unknown.

Here, we use *T. castaneum*, a genetically tractable coleopteran model, in which chitin biosynthesis is well-studied, to test whether Fabp-dependent regulation of Chs1 is conserved and functionally relevant for cuticle formation. Through comparative genomics, isoform-resolved expression profiling, RNAi-based functional analyses, immunohistochemistry, and biophysical characterization of the elytral cuticle, we identify TcFabp8 as an essential regulator of TcChs1-dependent chitin deposition and nanoscale cuticle organization in pupal elytra.

## 2. Results

### 2.1. Identification of *TcFABP* genes and splice variants in *T. castaneum*

A previous study performed in *D. melanogaster* identified a physical association between a Fabp, encoded by a single gene, and the chitin synthase 1, Krotzkopf verkehrt (Kkv) [17]. To test whether a comparable relationship might also exist in non-dipteran species, we identified Fabp homologs in the coleopteran pest model *T. castaneum*. We used *Drosophila* Fabp (DmFabp) sequence as a qBLASTp query and identified three putative Fabp homologs in the *T. castaneum* genome (Fig. 1A), which we named TcFabp2, TcFabp4, and TcFabp8 according to the chromosomal linkage group of their genes (Fig. 1B).

**Figure 1.**
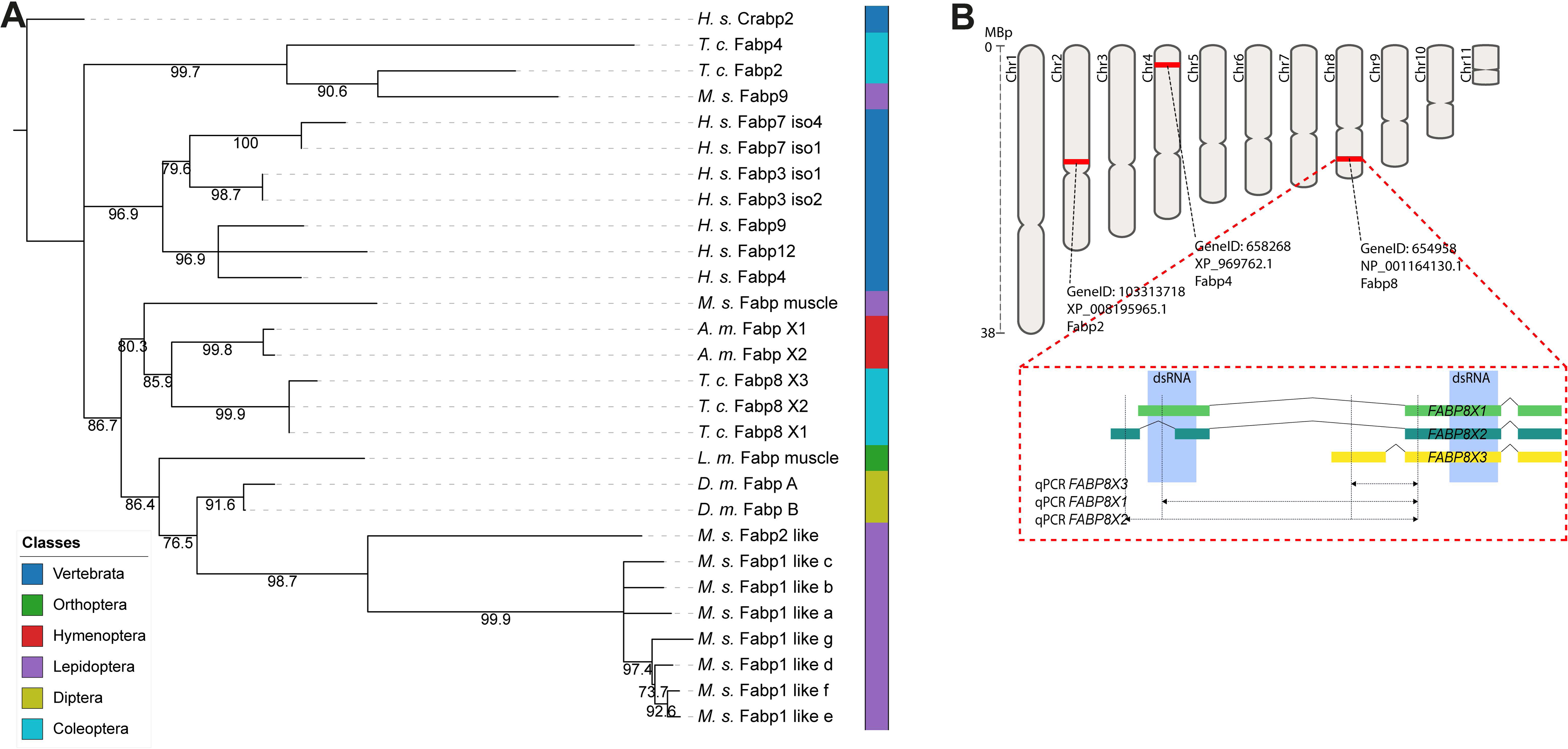
Phylogenetic analysis of Fabp family members and gene structure of *TcFABP8* isoforms. **(A)** Phylogenetic tree of Fabp family proteins from arthropod and vertebrate species. Major taxonomic groups are indicated by color. The *T. castaneum* sequences cluster within the broader Fabp family and were used to identify candidate orthologs for further functional analysis. Human Crabp2 was used as an outgroup. Branches with low bootstrap support were collapsed. (*H.s. Homo sapiens, T.c. Tribolium castaneum, M.s. Manduca sexta, L.m. Locusta migratoria, A.m. Apis mellifera, D.m. Drosophila melanogaster*) **(B)** Schematic overview of the chromosomal positions of three *T. castaneum FABP* loci (*TcFABP2*, *TcFABP4*, and *TcFABP8*) and detailed exon organization of the three *TcFABP8* isoforms. The upper panel shows the relative chromosomal positions of the three *FABP* loci. The lower panel shows an enlarged view of the exon structure of the three *TcFABP8* variants. Blue boxes indicate the dsRNA target regions used for RNA interference, and dashed lines indicate the isoform-specific qPCR amplicons. *TcFABP8X1* and *TcFABP8X2* differ in their 5′ region, whereas *TcFABP8X3* contains an alternative first exon.

To figure out which of these homologs might associate with TcChs1, we immunoprecipitated TcChs1 from *T. castaneum* extracts using a validated anti-Chs1 antibody directed to a C-terminal peptide and analyzed co-enriched proteins by mass spectrometry after filtering against anti-GFP controls. TcFabp8 (TC015275) was reproducibly detected in TcChs1 pull-downs above background (Fig. 2C). Although TcFabp8 did not pass the formal significance threshold in the statistical enrichment analysis, its repeated recovery in independent Chs1 pull-downs, together with the absence or weaker representation of the other TcFabp homologs, identified TcFabp8 as the most promising Fabp candidate for a potential association with TcChs1 and for subsequent functional analysis.

**Figure 2.**
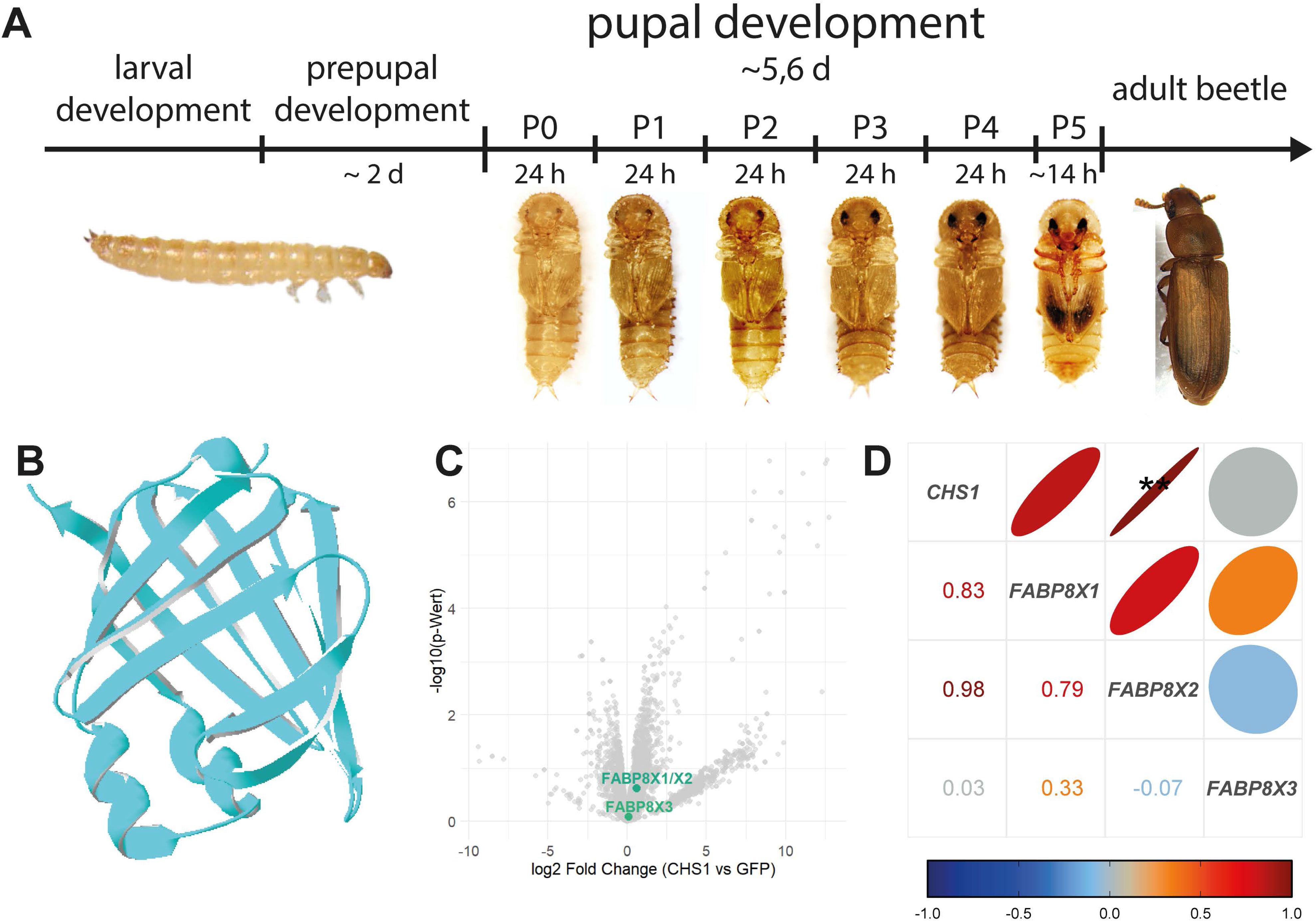
Developmental correlation of expression and identification of TcFabp8 as a candidate TcChs1-associated factor. **(A)** Schematic overview of *T. castaneum* development from larval to adult stages, including the prepupal period and pupal stages P0–P5. Representative images of the corresponding developmental stages are shown. The prepupal stage lasts approximately 2 days, followed by pupal development over approximately 5 to 6 days, with P0–P4 each representing 24 h and P5 approximately 14 h. **(B)** Predicted three-dimensional structure of TcFabp8, showing the characteristic Fabp fold. **(C)** Volcano plot of proteins identified in TcChs1 immunoprecipitation followed by mass spectrometry, comparing anti-TcChs1 pull-downs with GFP controls. TcFabp8X1/X2 and TcFabp8X3 are highlighted. The x-axis shows log2 fold change (TcChs1 vs GFP), and the y-axis shows the corresponding –log10(P) value. **(D)** Pearson correlation matrix showing the relationship between *TcCHS1* and *TcFABP8* isoform expression. Ellipses and color indicate the direction and strength of correlation, and the corresponding Pearson correlation coefficients are shown in the lower triangle. *TcCHS1* expression shows a strong positive correlation with *TcFABP8X1* and *TcFABP8X2*, whereas *TcFABP8X3* shows little or no correlation.

### 2.2. Phylogenetic analysis of Fabp proteins

To analyze TcFabp8 in a broader phylogenetic context, we examined its position among curated Fabp and Fabp-like proteins from representative vertebrate and invertebrate taxa, including selected arthropod lineages and human Fabp homologs. Moreover, we analyzed the exon-intron architecture of the *TcFABP8* locus and obtained indications for three splice variants by comparing genome and transcriptome data (see below). Multiple sequence alignment showed the expected conservation of the Fabp core domain across the dataset (Fig. S2). Maximum-likelihood reconstruction recovered several strongly supported local clusters, whereas deeper internal branching remained only partially resolved (nodes with bootstrap support < 70% were collapsed, Fig. 1A). Rooting with human cellular retinoic acid-binding protein 2 (Crabp2) as an external reference revealed that the three TcFabp8 splice variants form a tightly related, well-supported cluster within the broader Fabp/Fabp-like protein family. This TcFabp8 cluster grouped closely with hymenopteran Fabp sequences, whereas *Drosophila* Fabp proteins were placed separately together with *L. migratoria* Fabp and a distinct lepidopteran branch. In parallel, *Manduca sexta* displayed an expanded Fabp1-like subgroup, consistent with lineage-specific duplication and diversification. The two additional *T. castaneum* Fabp candidates identified in our BLAST search were placed outside the TcFabp8 cluster, consistent with more divergent Fabp/Fabp-like paralogs rather than members of the same local sequence group. Together, these data place TcFabp8 within the insect Fabp/Fabp-like protein family, while deeper ortholog relationships remain unresolved.

### 2.3. Analysis of *TcFABP8* splice variants

Database annotation identified three experimentally verified and two predicted TcFabp8 splice variants, each comprising three to four exons with an identical C-terminus but alternative N-terminal regions (Fig. 1B). Subsequent analyses focused on the three curated splice variants. Splice variants 1 and 2 encode the same core Fabp protein but differ in their N-terminal organization: *TcFABP8X1* contains an additional 5′ coding segment that gives rise to a predicted Sec/SPI signal peptide, whereas *TcFABP8X2* lacks this sequence. SignalP6.0 prediction indicates extracellular targeting for TcFabp8X1, whereas TcFabp8X2 and TcFabp8X3 are predicted to be cytosolic proteins (Fig. S3, Table S4). AlphaFold modeling confirmed that despite these N-terminal differences, all three isoforms share a canonical ten-stranded β-barrel core with a helix-turn-helix cap, consistent with a functional Fabp fold in each case (Fig. 2B, Fig. S1). To assess whether the expression of *TcFABP8* splice variants is developmentally correlated with *TcCHS1* expression, we profiled normalized transcript levels of *TcCHS1*, *TcFABP8X1*, *TcFABP8X2*, and *TcFABP8X3* in developing elytra at 24-hour intervals across pupal stages P0–P4, covering the complete pupal period of *T. castaneum* (Fig. S5). All three *TcFABP8* splice variants and *TcCHS1* showed their highest expression levels at P4, the stage of active cuticle deposition. Pearson correlation analysis revealed a strong positive association between *TcFABP8X2* and *TcCHS1* expression across the five pupal time points sampled (r = 0.98, p < 0.01; Fig. 2D), while *TcFABP8X1* showed a positive but non-significant trend (r = 0.83; Fig. 2D). *TcFABP8X3* expression was not correlated with *TcCHS1* expression (r = 0.03; Fig. 2D). The observed correlation of the expression of splice variant *TcFABP8X2* with *TcCHS1* may suggest a functional role of these proteins in cuticle formation during the active phase of elytral cuticle deposition.

### 2.4. Functional analyses of *TcFABP8* by RNAi

To test whether TcFabp8 is required for Chs1-dependent cuticle formation, we combined RNAi-mediated gene silencing with histological analysis of developing elytra at pupal stage P4 and phenotypic assessment of adult animals after ecdysis (developmental timepoints are shown in Fig. 2A). Knockdown of *TcFABP8* using a dsRNA targeting all splice variants resulted in a RNAi efficiency between 86 - 94% (Fig. S6, Table 1) and a loss of Fabp-immunoreactivity in immunohistochemistry (Fig. 4A). Monitoring the resulting phenotypes revealed that many pupae failed to shed the pupal cuticle (∼60%), whereas a smaller fraction emerged with incompletely matured and weakly pigmented elytra (∼20%; Fig. 3A). Importantly, these phenotypes could be reproduced by injecting a dsRNA targeting only splice variants *TcFABP8X1* and *TcFABP8X*2 at a RNAi efficiency of about 98% (Fig. 3D, Table 1), whereas the expression of *TcFABP8X3* remained unaffected (Table 1). This finding suggests that the phenotypic effects of silencing *TcFABP8* result from the activities of *TcFABP8X1* and/or *TcFABP8X*2. Isoform-specific RNAi targeting *TcFABP8X1* or *TcFABP8X2* individually was not feasible due to the high sequence identity within their shared coding region, which prevents a selective knockdown.

**Figure 3.**
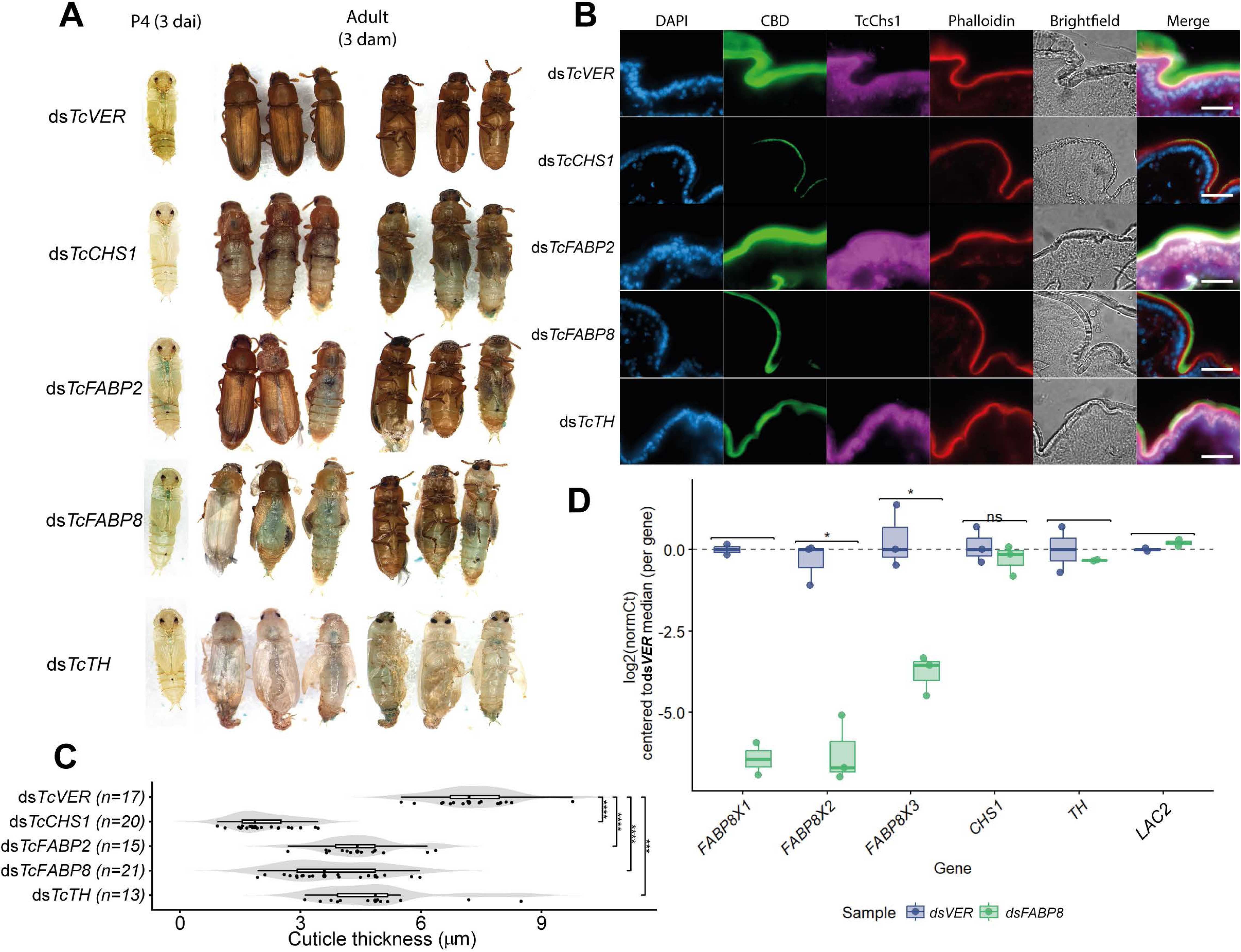
ds*TcFABP8* causes pigmentation defects, reduced Chs1 immunosignal, decreased cuticle thickness, and efficient depletion of Fabp8. **(A)** Representative phenotypes after RNAi of *vermilion* (*dsTcVER*), *TcCHS1* (ds*TcCHS1*), *TcFABP2* (ds*TcFABP2*), *TcFABP8* (ds*TcFABP8*), and *tyrosine hydroxylase* (ds*TcTH*). Compared with ds*TcVER*, ds*TcFABP8-*injected beetle showed reduced pigmentation and defects in adult cuticle formation. (dai = days after injection, dam = days after molting). **(B)** Immunohistochemistry of 15-µm cryosections of P4 elytra. Nuclei stained with DAPI (blue), chitin labeled with CBD-FITC (green), Chs1 detected with anti-Chs1 and goat anti-rabbit Cy3 (magenta), and actin visualized with phalloidin-ATTO633 (red). Brightfield images and merged channels are shown. In ds*TcVER* controls, Chs1 signal was detected at the apical cuticle-forming region, whereas this signal was strongly reduced in ds*TcFABP8-* and ds*TcCHS1*-injected pupae. Scale bars, 20 µm. **(C)** Quantification of cuticle thickness in P4 elytral cryosections following RNAi treatment. Violin plots show the distribution of values, with boxplots indicating the median and interquartile range. Each point represents one biological replicate, obtained by averaging three independent measurements per individual. Cuticle thickness was significantly reduced in all RNAi conditions compared to *dsTcVER*, with the strongest reduction observed after silencing *TcCHS1*, followed by *TcFABP8*, whereas *TcFABP2* and *TcTH* displayed intermediate effects. Differences between *TcFABP2* and *TcFABP8* were not statistically significant. Statistical significance was assessed using a Kruskal-Wallis test followed by pairwise Wilcoxon rank-sum tests with Benjamini-Hochberg correction for multiple comparisons. n = [13–21] animals/condition. **(D)** qPCR analysis of gene expression after RNAi of *TcFABP8*. Expression values are shown as log2(normCt) centered to the ds*TcVER* median for each gene. Knockdown of *FABP8* resulted in strong depletion of *FABP8X1*, *FABP8X2*, and *FABP8X3* transcripts, whereas expression of *TcCHS1*, *TcTH*, and *TcLAC2* was not markedly altered compared with controls. Boxes indicate the median and interquartile range; points represent individual biological replicates. Significance levels are indicated as follows: P < 0.05 (*), P < 0.01 (**), P < 0.001 (***), and P < 0.0001 (****).

**Figure 4.**
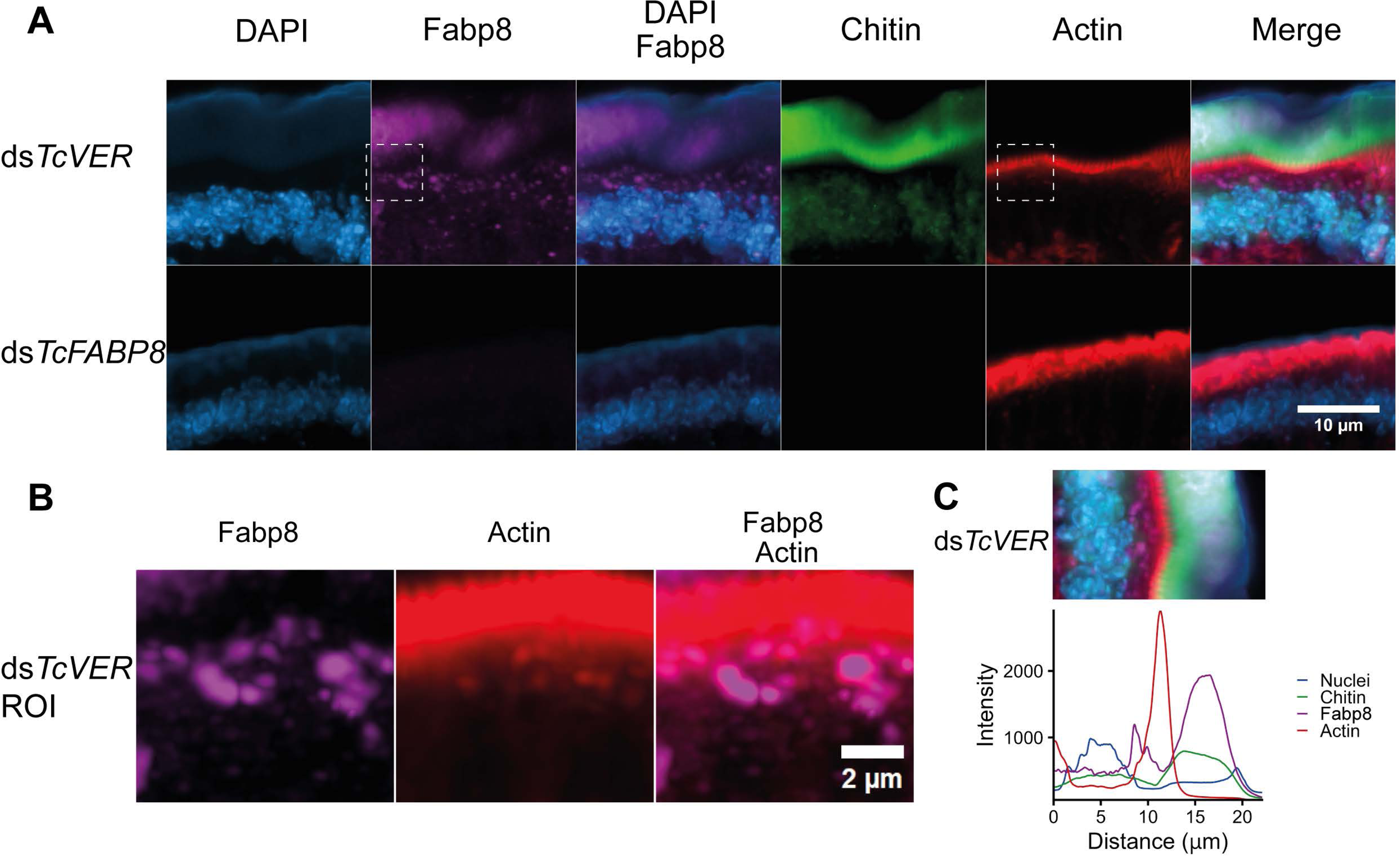
TcFabp8 localizes to the epidermis and is also detected within the cuticle of P4 elytra. **(A)** Immunohistochemical analysis of 15-µm cryosections of P4 elytra showing the localization of TcFabp8 relative to nuclei, chitin, and actin. Nuclei are stained with DAPI (blue), chitin was labeled with CBD-FITC (green), TcFabp8 was detected in magenta, and actin was visualized with phalloidin (red). Single channels and merged images are shown. TcFabp8 signal was detected in epidermal cells and in the cuticle. Scale bar, 10 µm. **(B)** Enlarged view of the boxed region in (A), highlighting TcFabp8-positive punctate structures in the apical epidermal region below the actin-rich zone and additional signal within the cuticle. Scale bar, 2 µm. **(C)** Representative line profile across the epidermis and cuticle showing the relative signal intensities of nuclei, chitin, TcFabp8, and actin along the indicated axis. The profile illustrates the spatial separation of the nuclear layer, the apical actin-rich region, and the chitin-rich cuticle, with TcFabp8 signals detected both below and within the cuticle region.

Overall, the resulting molting defects resembled the phenotypes observed after silencing *TcCHS1,* suggesting that chitin synthesis and cuticle formation may be impaired by RNAi targeting *TcFABP8*. To test this hypothesis, P4 cryosections from control and dsRNA-injected pupae were stained with primary antibodies against the C-terminus of Chs1 and secondary antibodies coupled to the fluorescent dye Cy3 to monitor its cellular distribution, a FITC-labeled chitin-binding domain (CBD) probe to visualize chitin deposition, DAPI to label nuclei, and phalloidin-ATTO633 to decorate F-actin filaments. As a positive control for RNAi, we used dsRNA targeting *vermilion* (ds*TcVER*), which produces a characteristic eye pigmentation defect but does not interfere with development, cuticle morphogenesis or chitin deposition.

Across all treatments, a strong actin signal was detected at the apical site of the epidermal epithelium, corresponding to the cortical actin network (Fig. 3B). This organization remained unaffected by any knockdown, indicating preserved epithelial integrity. In ds*TcVER* controls, TcChs1 immunoreactivity resulted in a strong signal at the apical epithelial surface, coinciding with robust chitin staining and the formation of a cuticle with a median thickness of ∼ 7.5 µm (Fig. 3B, C). Silencing *TcCHS1* (RNAi efficiency 80%; Fig. S6) abolished detectable TcChs1 immunoreactivity and led to thinner cuticle of approximately 2 µm, which was weakly chitinized, resulting in fully penetrant pharate-adult lethality.

Silencing *TcFABP8* caused a strong loss of TcChs1 immunoreactivity and a reduced chitin staining in the cuticle which appeared thinner (see below), similar to ds*TcCHS1* injected pupae. Thus, TcFabp8 depletion strongly disrupts the normal Chs1-associated chitin deposition pattern in developing elytra. Compared with RNAi for *TcFABP8*, RNAi for *TcFABP2* produced a milder developmental phenotype. However, TcChs1 immunoreactivity was not abolished in ds*TcFABP2* injected pupae. Instead, it appeared stronger and more broadly distributed across the epithelium than in controls, while chitin staining remained comparatively robust and cuticle thickness was less strongly affected (∼5 µm median; Fig. 3B,C). Thus, not all *Tribolium* Fabp/Fabp-like proteins phenocopy the strong Chs1-associated cuticle defect observed after TcFabp8 depletion.

Quantitative analysis of cuticle thickness across conditions confirmed these observations (Fig. 3C). Thickness differed significantly between RNAi treatments (Kruskal–Wallis test, p < 0.001). All tested *TcFABP8* knockdowns showed a significant reduction compared to *dsTcVER* controls (pairwise Wilcoxon tests with Benjamini– Hochberg correction, p < 0.001 for all comparisons). RNAi for *TcCHS1* resulted in the most severe reduction, followed by RNAi for *TcFABP8.* In contrast, RNAi for *TcFABP2*, as well as for *TcTH,* which encodes a tyrosine hydroxylase required for sclerotization, displayed intermediate phenotypes. Differences between *TcFABP2* and *TcFABP8* were not statistically significant, consistent with partially overlapping but distinct effects on cuticle formation. Analyses using a linear mixed-effects model on raw measurements yielded consistent results, confirming that the observed differences are robust and not driven by inter-individual variation.

### 2.5. Histochemical analysis of the elytral integument of dsRNA-injected pupae

To assess the localization of TcFabp8 and TcChs1, we performed immunohistochemistry and Airyscan CLSM microscopy using a commercially available antibody against human FABP4 (hFABP4) and anti-TcChs1 antibodies. Antibody specificity was supported by the loss of immunoreactivity after ds*TcFABP8 or* ds*TcCHS1* injection (Fig. 3B, 4A & 5A). In control elytra, TcFabp8 immunoreactivity appeared as discrete vesicle-like puncta in the subapical cytoplasm beneath the cortical actin network and was also detectable in the extracellular cuticle matrix (Fig. 4A-C). The extracellular TcFabp8 signal is consistent with the presence of an N-terminal signal peptide in TcFabp8X1. TcChs1 showed a subapical vesicular pool in addition to its strong membrane-associated apical signal. Together, these staining patterns indicate that TcFabp8 and TcChs1 are both present in subapical vesicle-like puncta beneath the apical deposition zone. Direct co-staining further supports recurrent partial spatial overlap of both proteins in a subset of these structures, consistent with a partially shared subapical compartment rather than complete co-residence in all vesicle-like puncta.

To directly assess the spatial relationship between TcFabp8 and TcChs1, we performed co-staining using ATTO647N-labeled anti-TcChs1 antibody together with indirect FITC detection of TcFabp8. In ds*TcVER* control sections, overlay images revealed recurrent local overlap of TcFabp8 and TcChs1 signals in discrete subapical vesicle-like puncta (Fig. 5A, 5B). This overlap was partial rather than complete, as single-positive signal regions were also observed and the two channels were not fully congruent. Line-profile analysis across the indicated subapical region showed corresponding local intensity maxima in both channels at several positions, supporting local spatial overlap of TcFabp8 and TcChs1 in a partially shared vesicle-like compartment beneath the cuticle-deposition zone (Fig. 5C).

**Figure 5.**
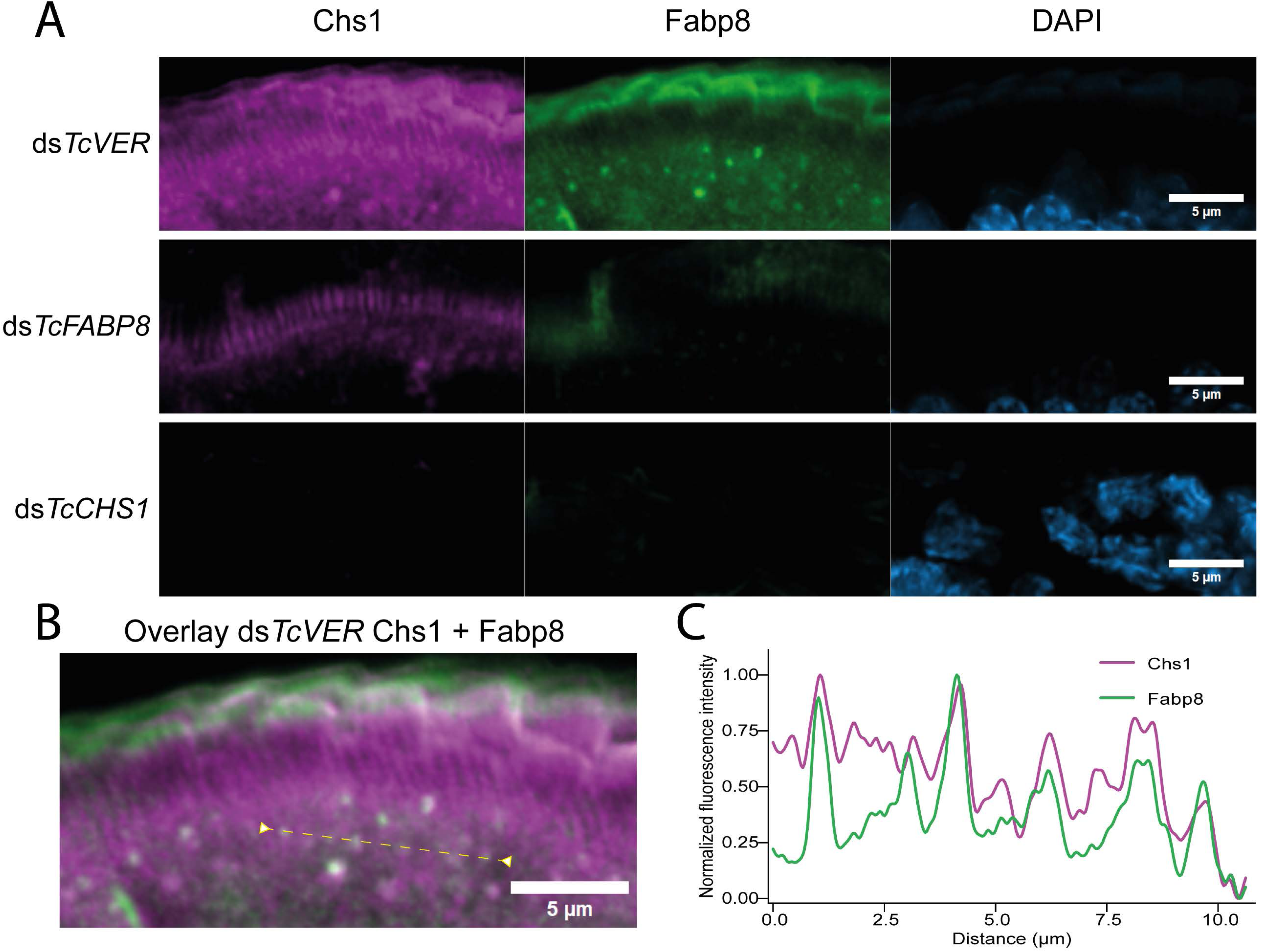
TcFabp8 and TcChs1 show partial spatial overlap in subapical vesicle-like puncta of pupal elytra. **(A)** Representative confocal images of P4 pupal elytral sections from ds*TcVER* control, ds*TcFABP8*, and ds*TcCHS1* animals stained for Chs1, Fabp8, and DAPI. In ds*TcVER* controls, Chs1 and Fabp8 signals were detected in the apical/subapical region of the epidermis and cuticle. Fabp8 signal was reduced after ds*TcFABP8* treatment, whereas Chs1 signal was strongly reduced after ds*TcCHS1* treatment, supporting antibody specificity. DAPI labels nuclei. **(B)** Overlay of Chs1 and Fabp8 signals in a ds*TcVER* section. The dashed line indicates the region used for line-profile analysis. **(C)** Normalized fluorescence intensity profile along the line shown in **(B)**. Corresponding local intensity maxima in the Chs1 and Fabp8 channels indicate recurrent partial spatial overlap in subapical vesicle-like puncta. Images were cropped for presentation. Linear brightness and contrast adjustments were applied to entire images or entire color channels only and were kept identical for images directly compared within each staining channel. No local adjustments, background removal, or signal-obscuring processing was applied. Scale bars, 5 µm.

We next asked whether these TcFabp8-positive subapical structures were associated with lipid-rich compartments during cuticle formation. For this purpose, pupal elytra cryosections were co-stained with CFW as a chitin marker, anti-hFABP4 antibody to localize TcFabp8, and Nile Red, a solvatochromic lipid dye, which was imaged in two spectral windows (488 nm excitation, 570 - 620 nm emission; 561 nm excitation, 620 - 700 nm emission) to capture differences in Nile Red fluorescence associated with local lipid microenvironments. [27, 28] In ds*TcVER* pupae, CFW staining confirmed chitin deposition in both the pre-existing pupal cuticle and the newly forming adult cuticle. TcFabp8 immunoreactivity was detected in the epidermal cell layer underneath the forming adult cuticle and in discrete subapical punctate structures, consistent with our earlier observations (Fig. 6A-C).

**Figure 6.**
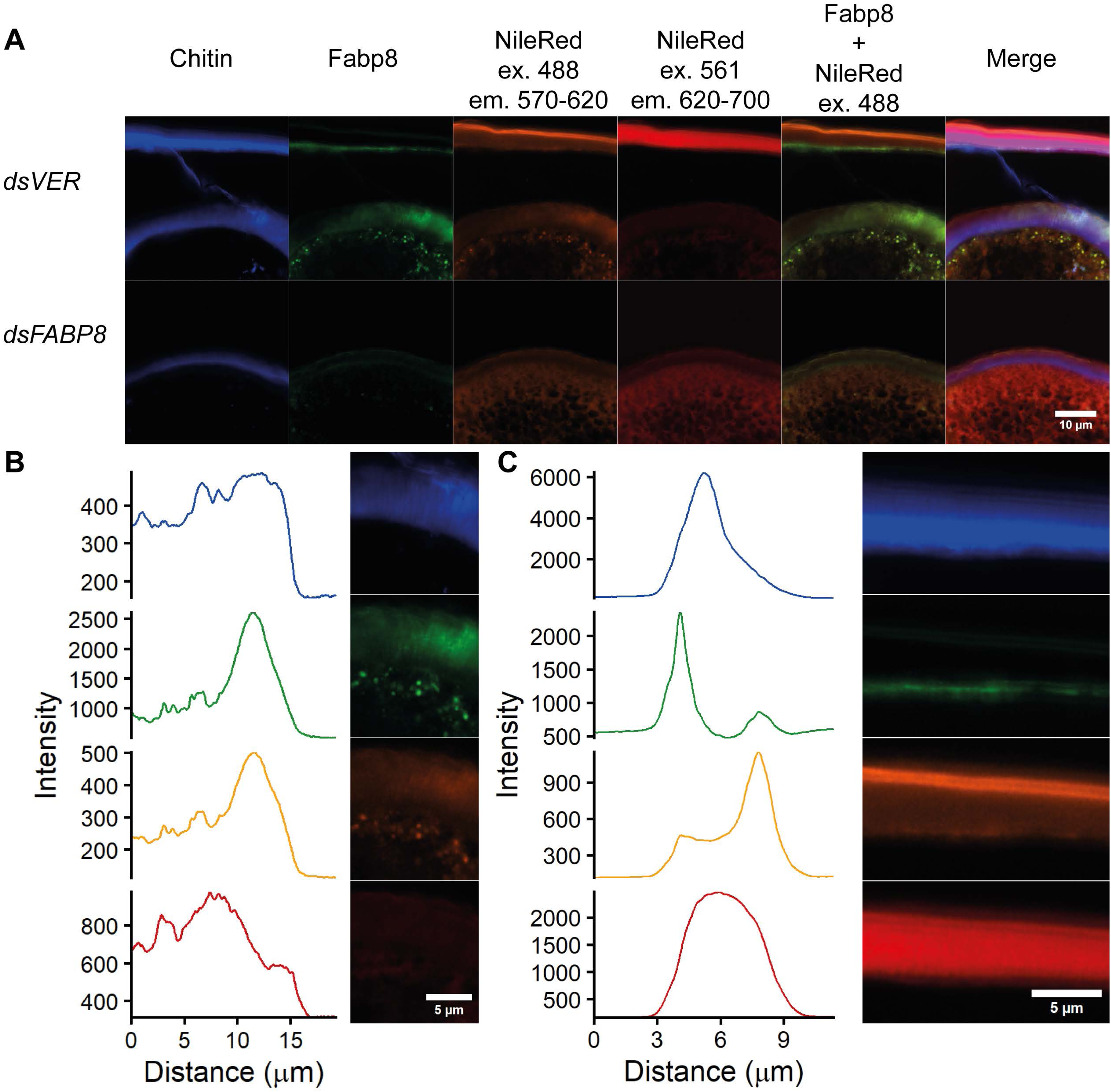
Nile Red reveals distinct lipid environments associated with TcFabp8 in pupal and developing adult elytra. **(A)** Fluorescence analysis of 15 µm elytral cryosections stained with Calcofluor White (CFW; chitin, blue), an anti-human FABP4 antibody detecting TcFabp8 (green), and Nile Red. Nile Red emission following 488-nm excitation is shown as the polar lipid channel (orange), whereas emission following 561-nm excitation is shown as the neutral lipid channel (red). Control ds*TcVER* pupae (upper row) show characteristic Nile Red distributions in pupal and newly forming adult cuticle. In pupal cuticle, the polar lipid signal is restricted to thin outer layers, whereas the neutral lipid signal is distributed more broadly. In adult cuticle, the polar lipid signal occurs within the cuticle and partially overlaps with TcFabp8. Following ds*TcFABP8* treatment (lower row), vesicular and cuticle-associated Nile Red signals are lost, while diffuse or accumulated intracellular signals predominate in both channels. Scale bar, 10 µm. **(B)** Newly forming adult cuticle. Left, representative line profile across the epidermis and cuticle showing CFW, TcFabp8, and polar and neutral Nile Red signals. Right, magnified view illustrating the spatial relationship between TcFabp8 and Nile Red within the cuticle. Scale bar, 5 µm. **(C)** Pupal cuticle. Left, representative line profile across the epidermis and cuticle. Right, corresponding magnified view. The polar lipid signal is confined to discrete outer layers, whereas the neutral lipid signal is distributed more broadly across the cuticle.

Nile Red fluorescence revealed distinct lipid distributions depending on the spectral window. In the Nile red channel acquired at 488 nm excitation (570 - 620 nm emission), the old pupal cuticle displayed a signal restricted to two narrow layers near the outer surface, whereas the forming adult cuticle exhibited a diffuse signal that overlapped spatially with TcFabp8 positive regions (Fig. 6A). High-magnification analysis confirmed that subapical TcFabp8 positive puncta partially overlaps with Nile Red fluorescence in this channel. Line profile analysis across the epidermis and forming cuticle revealed co-aligned signal peaks of TcFabp8 and polar Nile Red fluorescence in the subapical region, with CFW signal increasing more distally within the forming cuticle (Fig. 6B). In the other Nile red channel at 561 nm excitation (620 - 700 nm emission), the old pupal cuticle displayed strong and broadly distributed fluorescence, with an additional thin layer between the two outer polar lipid layers (Fig. 6C). In contrast, the forming adult cuticle showed only weak neutral lipid signal in this channel (Fig. 6B). Taken together, these data indicate that the TcFabp8-positive subapical puncta are associated with hydrophobic, lipid-rich vesicle-like structures and spatially precede chitin deposition, consistent with a role for TcFabp8 in organizing a hydrophobic, lipid-rich compartment at the site of cuticle formation. RNAi-mediated depletion of *TcFABP8*X1/2 resulted in a loss of subapical TcFabp8-positive signals and a marked redistribution of lipids: Nile Red fluorescence became strongly increased markedly and diffusely distributed throughout the epidermal cells in both spectral windows. Concomitantly, formation of the adult cuticle was abolished, as indicated by the absence of newly deposited CFW positive material. These findings indicate that TcFabp8 is required for the organization and transport of hydrophobic, lipid-rich vesicle-like structures to distinct sites within the apical cuticle.

### 2.6. Structural analysis of the cuticle nanoscale organization

Following *TcFABP8* RNAi, which leads to reduced cuticle thickness and loss of detectable Chs1 immunoreactivity at pupal stage P4, we asked whether these defects extend to the nanoscale organization of remaining chitin in the adult elytral cuticle using synchrotron micro-focused X-ray diffraction (XRD).

Elytra from ds*TcVER* controls displayed the characteristic diffraction pattern of α-chitin, including a fiber correlation peak at q ≈ 1.37 nm⁻¹ and well-resolved (020), (110), (013), and (120) reflections (Fig. 7A - B; [29–33]. The persistence of the characteristic α-chitin-associated scattering features, including reflections in the 020-, 110/120-, and 013- associated q-regions, supports retention of α-chitin-related organization. After scaled Kapton-background subtraction, peak fitting was performed separately for the low-q correlation peak and the WAXS region (Fig. S7, Table S5-S7). The low-q correlation peak was consistently shifted to lower q values in ds*TcFABP8* elytra compared with the paired ds*TcVER* controls (Table S6). The fitted peak center decreased from 1.3495 to 1.3303 nm⁻¹ in frame A, from 1.3514 to 1.3367 nm⁻¹ in frame B, and from 1.3519 to 1.3055 nm⁻¹ in frame C. This corresponds to an increase in the associated real-space correlation distance from approximately 4.65 nm in ds*TcVER* controls to approximately 4.72, 4.70, and 4.81 nm in ds*TcFABP8* elytra, respectively (Table S5). The fitted width of the low-q peak was also higher in all ds*TcFABP8* samples, although the effect was strongest in frame C (Table S5).

**Figure 7.**
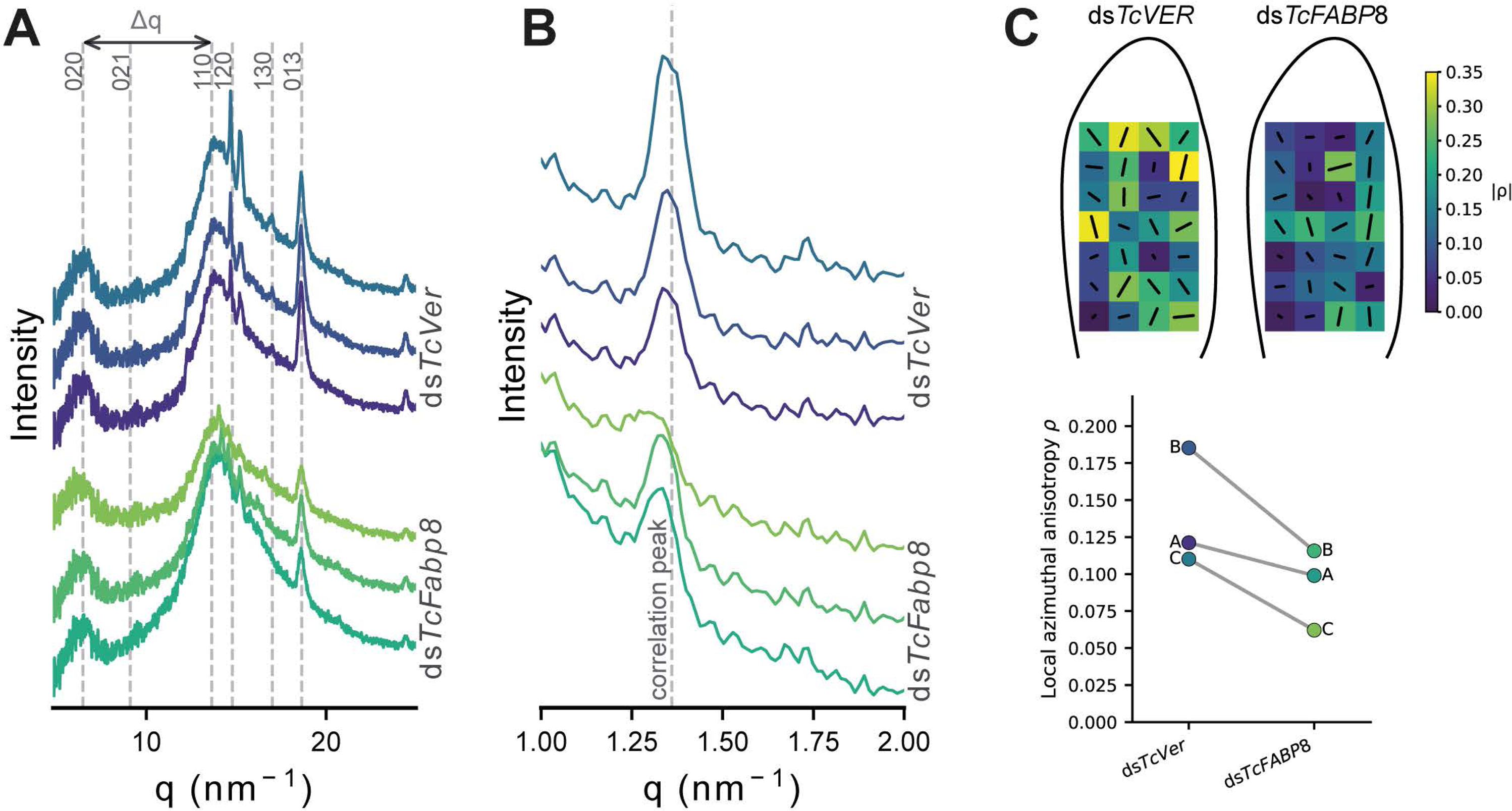
X-ray diffraction analysis of α-chitin organization in elytra from pupae injected with dsTcVER and dsTcFABP8. **(A)** One-dimensional X-ray diffraction (XRD) profiles of intact elytra from ds*TcVER-* injected controls and ds*TcFABP8*-injected pupae over a q-range of 3–25 nm⁻¹. Profiles are vertically offset for clarity but plotted without intensity normalization or rescaling. Vertical dashed lines indicate characteristic α-chitin reflections [(020), (021), (110), (120), (130), and (013)]. The spacing between the (110) and (020) reflections (Δ *q*_(110)–(020)_) is marked by a double-headed arrow. Data from three biological replicates (fr. A–C) are shown for each condition. **(B)** Enlarged low-q region (1–2 nm⁻¹) of the same stacked XRD profiles, highlighting the lamellar fiber correlation peak of α- chitin at q ≈ 1.37 nm⁻¹. **(C)** Schematic representation of elytra indicating the scanned regions for ds*TcVER* and ds*TcFABP8* treated samples. For each genotype, a representative vector map summarizes pixel-wise scattering orientations. **(D)** Each point represents the median of local scan positions within one paired elytron scan; lines connect ds*TcVER* and ds*TcFABP8* samples measured in the same frame.

Fits of the WAXS region confirmed the persistence of chitin-associated scattering features in ds*TcFABP8* elytra. The 18.6 nm⁻¹ component, corresponding to the α-chitin-associated high-q reflection, remained centered at a similar q position in both treatments, whereas its fitted amplitude tended to be lower and its fitted width higher in ds*TcFABP8* samples (Fig. S7, Table S7). Other WAXS components showed treatment-dependent changes in fitted width and amplitude, but these were not interpreted as individual quantitative changes in crystalline volume fraction because several components represent broad or overlapping peak envelopes and are sensitive to background modeling. Together, the scaled-background fits support preservation of α-chitin-associated scattering features, while indicating altered low-q correlation spacing and reduced structural coherence or increased heterogeneity of the residual cuticular scaffold after TcFabp8 depletion.

To assess the projected nanoscale organization of the low-q correlation signal, we quantified local detector-plane azimuthal anisotropy in the q = 1.30–1.40 nm⁻¹ range (Fig. 7C). Across all three paired scans, ds*TcFABP8* elytra showed lower median local ρ values than the corresponding ds*TcVER* controls measured in the same frame (Fig. 7D). Median ρ decreased from 0.121 to 0.099 in frame A, from 0.185 to 0.116 in frame B, and from 0.110 to 0.062 in frame C. Mean local ρ values showed the same trend, decreasing from 0.173 to 0.113 in frame A, from 0.187 to 0.118 in frame B, and from 0.160 to 0.072 in frame C (Fig. 7D, Table S8). Thus, TcFabp8 depletion consistently reduced the local detector-plane azimuthal anisotropy of the low-q correlation signal across all paired scans.

Together, these results indicate that TcFabp8 depletion does not alter α-chitin polymorph identity, but reduces the amount of ordered chitin fibrillar material and disrupts the higher-order nanoarchitecture of the cuticular scaffold, with less coherent fibril packing and more heterogeneous fibril alignment.

## 3. Discussion

The data provided in this study indicate that TcFabp8 is an important regulator of TcChs1-dependent cuticle formation in *T. castaneum* and possibly other insects. Chitin synthases are highly conserved across insect orders, and their regulation has been studied most extensively in *D. melanogaster* and *T. castaneum* [8]. A physical association between Fabp and the chitin synthase Kkv was previously reported in *D. melanogaster*, where Fabp depletion impaired wing chitin deposition [17]. We now show that a comparable relationship exists in a coleopteran species. TcFabp8 was reproducibly identified by mass spectrometry in co-immunoprecipitates generated with TcChs1 antibodies, and its RNAi-mediated depletion caused strong reduction of apical Chs1 immunoreactivity, diminished chitin incorporation, reduced cuticle thickness, and high-penetrance molting failure. Notably, α-chitin identity was preserved in RNAi survivors, indicating that TcFabp8 is not required for chitin allomorph specification, but rather for efficient Chs1-dependent chitin deposition.

This view is consistent with emerging evidence that Fabp-dependent lipid handling contributes to developmentally critical processes in other arthropods, although through context-dependent mechanisms. In *Locusta migratoria*, juvenile hormone analogue exposure disrupts hormone and lipid-metabolic pathways and strongly suppresses Mg-

FABP expression; RNAi-mediated Mg-FABP depletion reduces triglyceride levels and lipid droplet accumulation in the fat body, linking insect Fabp function to developmental lipid homeostasis [25]. In the acarine ectoparasite *Dermanyssus gallinae*, DgFabp localizes to several tissues including the cuticle, and its knockdown increases mortality, reduces deutonymphal molting, and impairs male mating performance [34]. In *B. mori* Fabp2/3 expressed in the midgut were linked with a lipid homestatic imbalance leading to a pigmentation-related phenotypic output affecting the cuticle [26]. However, these studies do not establish a Chs1-dependent mechanism, but they support the broader idea that Fabp-mediated lipid handling can influence molting, tissue maturation, and pest-relevant developmental outputs across arthropods.

TcFabp8 appears to act at the post-translational level, playing a role in proper Chs1 localization and stability. RNAi-mediated depletion of TcFabp8 did not reduce Chs1 transcript abundance, whereas immunohistochemistry revealed a near-complete loss of detectable apical Chs1 immunoreactivity (Fig. 3B, 3D). This finding suggests that TcFabp8 is required for the proper spatial distribution of Chs1, thereby promoting TcChs1 retention and/or apparent protein stability at the apical membrane. The reduced cuticle thickness that mirrors the loss of Chs1 activity probably reflects this effect. In control elytra, TcFabp8 is localized to subapical puncta beneath the cortical actin network, a compartment consistent with recycling endosomes and exocytic vesicle pools that supply apically directed cargo [35]. Notably, Chs1 immune signals displayed a similar subapical vesicular-like staining in addition to its localization at the apical membrane, consistent with a regulated vesicular delivery of Chs1 containing vesicles to the deposition zone – a trafficking mode previously described for Kkv in *Drosophila* and linked to cytoskeletal and calcium-dependent regulatory cues [18]. Importantly, direct co-staining using ATTO647N-labeled anti-TcChs1 together with indirect FITC detection of TcFabp8 revealed recurrent partial spatial overlap of both signals in discrete subapical vesicle-like puncta. This overlap was not complete, and single-positive regions were also observed, indicating that TcFabp8 and TcChs1 are not constitutively present in an identical compartment. Rather, the data support a model in which TcFabp8 associates with a partially shared subapical vesicle-like compartment that promotes TcChs1 retention, stabilization, or delivery to the apical cuticle- deposition machinery. Fabps are known to modulate membrane lipid composition and influence vesicle dynamics through their capacity to shuttle fatty acids between intracellular compartments [36], raising the possibility that TcFabp8 additionally shapes the lipid environment required for efficient Chs1 vesicle formation or fusion at the apical membrane.

The scaled-background XRD and SAXS analyses refine the hierarchical level at which TcFabp8 affects cuticle formation. Insect cuticle assembly proceeds through successive organizational levels, from chitin polymer deposition to fibril bundling, lamellar stacking, and final sclerotization [19]. In TcFabp8-depleted elytra, recognizable α-chitin-associated scattering features were retained, and the high-q component near 18.6 nm⁻¹ remained positionally stable, supporting the conclusion that TcFabp8 depletion does not abolish α-chitin-associated organization (Fig. 7). Instead, the most robust structural changes were observed in the low-q correlation peak, which shifted consistently to lower q-values and showed increased fitted width in ds*TcFABP8* samples, corresponding to an increased correlation distance and a broader distribution of low-q structural correlations. This argues against a simple change in α-chitin allomorph identity and instead points to altered mesoscale packing or scaffold organization of the residual cuticular matrix. In the WAXS region, the 18.6 nm⁻¹ α- chitin-associated component tended to show reduced fitted amplitude and increased width, whereas other peak amplitudes were not consistently reduced across paired scans. Together with the reduced detector-plane azimuthal anisotropy of the low-q correlation signal, these data suggests that TcFabp8 affects the projected local organization and coherence of the residual cuticular scaffold. Because several WAXS fit components represent broad or overlapping peak envelopes, these results should not be interpreted as a direct quantitative measure of crystalline volume fraction, absolute three-dimensional fibril disorder, or unit-cell-level lattice changes. Fabps are broadly understood as intracellular lipid chaperones that regulate the availability and intracellular trafficking of long-chain fatty acids and other hydrophobic ligands [36]. This interpretation is supported by recent functional work in *L. migratoria*, where Mg-Fabp depletion reduced fat-body lipid droplet accumulation, indicating that insect Fabps can control intracellular lipid distribution and storage *in vivo* [25].

The canonical β-barrel fold and helix-turn-helix structure of the TcFabp8 core domain, predicted by AlphaFold modeling (Fig. S1), is consistent with this function and places TcFabp8 structurally within the well-characterized Fabp family despite the absence of direct ligand-binding data. Beyond their classical role as cytosolic carriers, Fabps have been shown to interact directly with membranes and to influence local lipid composition, thereby modulating membrane curvature, fluidity, and vesicle budding [37].

The insect cuticle is known to contain spatially segregated lipid layers with distinct compositions [19, 38]. Dual-channel Nile Red imaging in two spectral windows was therefore used to discriminate between different lipid microenvironments, based on the solvatochromic properties of the dye [27, 28]. In control pupae, Nile Red-channel signal revealed a subapical population of lipid-associated puncta that partially colocalizes with TcFabp8-positive puncta, whereas signal in the red-shifted detection window is largely absent (Fig. 6). The red channel subapical signal was lost upon RNAi-mediated depletion of TcFabp8; instead, a more diffuse intracellular fluorescence pattern was observed in both channels (Fig. 6). In addition to this intracellular redistribution, TcFabp8 depletion was associated with a shift in the spatial distribution of lipid-associated signals between the epidermal cell layer and the cuticle (Fig. 6). In controls, Nile Red fluorescence was detectable in both compartments, with signals in the polar lipid channel present in both cuticle and cellular layers, and signals in the neutral lipid channel predominantly enriched in the cuticle (Fig. 6A). Upon RNAi to silence *TcFABP8*, lipid-associated fluorescence appeared relatively increased within the cellular layer and reduced in the cuticle across both detection channels, consistent with a redistribution of lipids from the extracellular matrix to intracellular compartments. This shift was more pronounced in the neutral lipid channel, suggesting that TcFabp8 may differentially affect the deposition or retention of distinct lipid classes and microenvironments within the cuticle. Notably, these changes occur without an apparent reduction of lipid-associated signal, indicating that TcFabp8 may influence lipid organization rather than lipid abundance [36, 39]. These observations are consistent with a model in which TcFabp8 organizes a subapical lipid environment, characterized by laterally segregated membrane domains, that supports the vesicle-associated delivery of Chs1 to the apical membrane and its functional deployment at the chitin deposition zone [18, 35, 40].

The transcription of different splice variants from the TcFabp8 suggests functional compartmentalization. TcFabp8X1 and TcFabp8X2 encode identical core Fabp domains but differ in their N-terminal organization, with a Sec/SPI signal peptide predicted only for TcFabp8X1 (Fig. S1), suggesting potential extracellular targeting. Signal peptide-dependent secretion of lipid-binding proteins into extracellular compartments has been described in both nematode and insect systems. In nematodes, Fabps carrying N-terminal signal peptides are secreted into the perivitelline fluid surrounding developing embryos [41]. More recently, a noncanonical α-helical Fabp, Obp44a, present in *Drosophila* glia was shown to be secreted via a signal peptide-dependent mechanism into the hemolymph, where it functions as an extracellular lipid chaperone [39]. These examples support the possibility of an extracellular TcFabp8X1 pool. Consistently, TcFabp8 immunoreactivity was detected in subapical cytoplasmic puncta and within the cuticle, and was abolished after ds*TcFABP8* treatment, confirming the specificity of immune signals. The developmental co-regulation of *TcFABP8X2* and *TcCHS1* expression at P4 further supports the functional relevance of the cytosolic isoform pool during active cuticle deposition. The strong positive correlation between *TcFABP8X2* and *TcCHS1* transcript levels across pupal development (r = 0.98) suggests that both the secreted and cytosolic TcFabp8 pools are transcriptionally primed in concert with the chitin biosynthetic machinery (Fig. 2D), consistent with a coordinated rather than independent deployment of the two isoform pools during cuticle assembly. The successful recombinant expression and purification of TcFabp8X2 (Fig. S4) confirms that the shared core Fabp domain folds independently from the presence or absence of a signal peptide. Together, these data support a two-pool model in which the TcFabp8 locus generates a cytosolic splice variant that may contribute to intracellular lipid transport to sites of Chs1 deposition, and a secreted splice variant that could function within the extracellular cuticular matrix, for instance by influencing the lipid environment during fibril assembly or consolidation. To our knowledge, this would represent the first description of a secreted Fabp within an insect cuticular matrix. Because of the shared core coding nucleotide sequence precluded a clean isoform-specific RNAi separation in the present study, the distinct functional roles of each pool remain to be established.

## 4. Conclusions

Taken together, our data establish TcFabp8 as a critical facilitator of TcChs1-dependent chitin deposition and cuticle maturation machinery in *T. castaneum* and extend the functional connection between lipid handling and chitin biosynthesis beyond the dipteran lineage [17]. The convergence of TcChs1 co-immunoprecipitation, RNAi phenotyping, quantitative histology, partial spatial overlap of TcFabp8 and TcChs1 in subapical vesicle-like puncta, lipid-associated Nile Red redistribution, and synchrotron diffraction analysis provides a multi-faceted picture in which TcFabp8 does not affect chitin’s allomorphic identity, but supports proper TcChs1 transport and localization, chitin accumulation, and higher-order organization of the cuticular scaffold. The architecture of the *TcFABP8* locus further suggests that these functions may be partitioned between cytosolic and putatively secreted protein pools [39, 41, 42], potentially allowing TcFabp8 to act both upstream of and within the growing cuticular matrix. Whether this two-pool organization represents a conserved feature of Fabp-dependent cuticle regulation across insects, and how TcFabp8 interfaces mechanistically with TcChs1 vesicle trafficking and apical secretion machinery remain important open questions. More broadly, our findings suggest that lipid-handling proteins of the Fabp family may be integral components of the cuticle biosynthetic apparatus rather than peripheral modulators [36, 37], a perspective that may impact future work on cuticle-related resistance mechanisms and Fabp-centered pest-control strategies in insects and other arthropods.

## 5. Materials and Methods

### 5.1. Insect rearing

All experiments were performed with the *T. castaneum* GA-1 strain [43], continuously maintained in the laboratory since 2009. Beetles were reared on whole-wheat flour under standard conditions (30 °C, 50% humidity, continuous darkness) as described previously [44]. Larvae, pupae, and adults were collected manually under a stereomicroscope.

### 5.2. Bioinformatic analyses

The amino acid sequences of *T. castaneum* Fabps were identified by NCBI qBLASTp using the *D. melanogaster* Fabp amino acid sequence (NP_001027180.1) as a query. Orthologs were selected based on ≥ 75% query coverage and ≥ 35% sequence identity. To examine phylogenetic relationships of TcFabps, additional homologs were retrieved using Entrez-BLAST from a manually curated species list. Multiple sequence alignments (MSA) were generated with MAFFT [45]and trimmed to optimize alignments. Phylogenetic trees were built using IQ-TREE [46] with automated model selection and bootstrap-method (maximum 1000 bootstrap replicates with bootstop). Trees were visualized in iTol [47] and finalized graphically in Adobe Illustrator. Structural modeling of *T. castaneum* Fabps and their isoforms was carried out in AlphaFold3 [48]. Prediction of signal peptides was performed with SignalP6.0 [49].

### 5.3. Co-immunoprecipitation and mass spectrometry analysis

Co-immunoprecipitation (Co-IP) was performed to identify candidate Chs1-associated proteins. Lysates from pools of 30 *T. castaneum* individuals per condition were subjected to Co-IP using Dynabeads M-270 Epoxy (Thermo Fisher Scientific) according to the manufacturer’s instructions. A custom rabbit polyclonal antibody raised against a C-terminal Chs1 peptide (SVASSIPAKD; ProteoGenix) was used for pulldown, with a commercial anti-GFP antibody serving as a negative control. Three biological replicates were performed for each condition. Eluted proteins were lyophilized and analyzed by mass spectrometry (Mass Spectrometry Core Facility, University of Osnabrück). Data were processed using the facility’s standard pipeline and resulting protein lists were comparatively screened in R for proteins reproducibly detected in Chs1 samples relative to controls.

### 5.4. RNA interference: dsRNA synthesis and injection

T7-overhang primers (Table S1) were designed to amplify selected cDNAs that were tested for unique genomic specificity of the resulting double-stranded RNA (dsRNA) using the dsRNAengineer web tool and synthesized by Invitrogen (USA). PCR amplification was performed with DreamTaq polymerase (ThermoFisher, USA) under the following conditions: 95 °C for 5 min; 35 cycles of 95 °C for 30 s, 60 °C for 30 s, and 72 °C for 60 s; followed by 72 °C for 2 min. Amplicons were purified using the Qiagen gel extraction kit (Qiagen, Germany). dsRNA was synthesized using the NEB HiScribe T7 High-Yield RNA Synthesis Kit (New England Biolabs, USA) as described previously [21].

For injections, borosilicate capillaries (*ScienceProducts*, Germany) were pulled using a Sutter P-97 micropipette puller (Sutter Instruments, USA). Settings for pulling were chosen based on the Sutter pipette cookbook and adapted slightly to result in consistent injection-needles. Approximately 200 nl of dsRNA solutions (400 ng dsRNA in injection buffer with diluted dye) were injected into P0 pupae at a maximum of 16 h after pupal molt using a micromanipulator and microinjection needle holder (Narishige, Japan). Each treatment group consisted of 20 pupae. After injection, animals were reared on whole wheat flour at 30 °C for 4 days prior to sampling for qPCR, histology or further reared for phenotypic analysis.

### 5.5. Quantitative PCR (qPCR)

For the determination of RNAi efficiency, three P4 pupae were pooled per replicate and treatment. For expression profiling, synchronized P0 pupae were maintained under standard conditions, and samples were collected every 24 h until adult emergence (stages P1–P4 and adult; P5 was dismissed). Elytra dissected from five individuals were pooled for each replicate. Total RNA was extracted using RNeasy Mini Kit (Qiagen, Germany) as recommended by the manufacturer. cDNA synthesis was performed with the RevertUP II cDNA Synthesis Kit (BiotechRabbit, Germany) at 52.5°C overnight, followed by heat inactivation at 95 °C for 10 min. cDNA concentrations were adjusted to 50 ng/µl. qPCR was carried out using a Mic qPCR Cycler (BMS, Australia) and Nippon SyGreen Master Mix (Nippon Genetics Europe, Germany). Each 20 µl reaction contained 100 ng cDNA, 500 nM primers, and 1x SyGreen master mix. Cycling conditions were: 95 °C for 5 min, followed by 40 cycles of 95 °C for 10 s, 60°C for 15 s, and 72 °C for 30 s. Specificity was confirmed by melt-curve analysis (72– 95 °C, Δ°C/T = 0,3 °C/s).

Relative expression was normalized to *TcRpS18* using the Pfaffl method [50], which was validated as a stable reference gene across tissues and stages [51]. Mean normalized expression values and RNAi efficiencies were calculated from three biological and three technical replicates.. Mean normalized expression values per pupal stage were calculated from three biological replicates for correlation analysis of developmental expression profiles. Pearson correlation coefficients for gene expression between *TcCHS1*, *TcFABP8X1*, *TcFABP8X2*, and *TcFABP8X3* were computed in R (version 4.2.2). Significance was assessed by two-tailed t-test. Asterisks indicate p < 0.05 (*), p < 0.01 (**), and p < 0.001 (***).

### 5.6. Cryosectioning and Immunohistochemistry

Elytra were dissected and fixed in 4% (w/v) formaldehyde (FA) in PBS for 1 h at room temperature (RT). Samples were cryoprotected in a graded sucrose series (12%, 15%, 18%, two times 20% for 20 min each at RT), embedded in OCT compound, and frozen in 2-butanol cooled with liquid nitrogen. Cryoblocks were trimmed and sectioned at 15 µm using a Leica cryostat at −19 °C.

Sections were mounted on poly-L-lysine-coated slides, dried for 5 min at 40 °C, and washed four times for 15 min in PBS and twice in PBS + 0.1 % Triton X-100 at RT. Primary antibodies (Table S2) diluted in PBS + 0.1% (v/v) Triton X-100 were applied overnight at 4 °C. Slides were washed as above and incubated with secondary antibodies (Table S2) and, where indicated, fluorescent dyes such as DAPI, CBD- FITC, and phalloidin-ATTO633 (Table S3) overnight at 4 °C. For lipid staining, Nile Red and Calcofluor White (CFW) were applied diluted in PBS (Table S3) after incubation with the secondary antibody. After final washing, sections were mounted in VectaShield and coverslips were sealed with Fixogum using ring reinforcers as spacers to prevent tissue compression.

### 5.7. Immunohistochemical co-staining of TcFabp8 and TcChs1

For colocalization analysis of TcFabp8 and TcChs1, P4 pupal elytra were dissected, fixed, cryoprotected, embedded, and sectioned as described above. Because both primary antibodies were raised in rabbit, a sequential staining strategy was used in which the anti-TcChs1 antibody was directly fluorescently labeled prior to co-staining.

The rabbit polyclonal anti-TcChs1 antibody was supplied in PBS containing 0.02% (w/v) sodium azide at a concentration of 1.01 mg/ml. The antibody was labeled with ATTO 647N NHS ester (ATTO-TEC, Siegen, Germany) using the Alexa Fluor™ Antibody Labeling Kit (Thermo Fisher Scientific, Cat. No. A88071) according to the manufacturer’s instructions, except that the supplied fluorophore was replaced by ATTO 647N NHS ester. A 10-fold molar excess of dye relative to antibody was used. After incubation for 15 min at RT, excess unreacted dye was removed using the supplied dye-removal spin columns. The labeled antibody was used directly or stored protected from light at 4 °C until use.

For co-staining, cryosections were washed and permeabilized as described above. Sections were first incubated overnight at 4 °C with anti-TcFabp8 antibody diluted in PBS containing 0.1% Triton X-100. After washing, TcFabp8 was detected with a goat anti-rabbit FITC-conjugated secondary antibody overnight at 4 °C. Sections were then washed four times in PBS at RT and fixed again in 4% formaldehyde in PBS for 10 min at RT to stabilize the antibody complexes. After two additional PBS washes and two washes in PBS containing 0.1% Triton X-100, sections were incubated overnight at 4°C with directly labeled anti-TcChs1–ATTO 647N diluted in PBS containing 0.1% Triton X-100. Nuclei were counterstained with DAPI. Sections were washed again in PBS and mounted as described before.

Negative controls were processed in parallel by omitting the primary antibodies. Signal specificity was further assessed using sections from ds*TcFABP8*- and ds*TcCHS1*- treated animals. Channels were acquired sequentially using non-overlapping detection windows to minimize spectral crosstalk and bleed-through. The absence of detectable signal in secondary-only controls and the reduction of antibody signals after the respective RNAi treatments supported staining specificity.

### 5.8. Confocal laser scanning microscopy and image processing

Fluorescence images were acquired with a Zeiss LSM 900 confocal laser scanning microscope (Zeiss, Germany) using 63x oil-immersion objective (NA=1.4). Excitation was performed with UV (405 nm), blue (488 nm), green (561 nm), and red (640 nm) diode lasers for DAPI, FITC, Cy3, Nile Red, and ATTO 633 or ATTO647N fluorophores, respectively. Channels were scanned sequentially to prevent bleed-through. For TcFabp8/TcChs1 co-staining, FITC was excited at 488 nm and detected at approximately 500–550 nm, whereas ATTO647N was excited at 640 nm and detected at approximately 650–700 nm. Nile Red fluorescence was recorded in two spectral windows to distinguish different lipid-associated environments. For the shorter-wavelength Nile Red channel, excitation was performed at 488 nm and emission was detected at approximately 570–620 nm. For the red-shifted Nile Red channel, excitation was performed at 561 nm and emission was detected at approximately 620– 700 nm. Images were collected in Airyscan super-resolution mode (pixel size 0.04 µm, line averaging 4, bit depth 16) and Airyscan-processed with constant Wiener filter strength to all compared images using ZEN Pro Blue (Zeiss, Germany) software. All images were postprocessed in Fiji (ImageJ) [52]. Brightness and contrast adjustments were applied uniformly across all samples. Final figure panels were assembled and annotated using Adobe Illustrator. Orientation and position are indicated relative to the beetle body axis.

For visualization of local spatial overlap between TcFabp8 and TcChs1, line profiles were drawn across the subapical region containing vesicle-like puncta. Fluorescence intensities of the TcFabp8 and TcChs1 channels were normalized independently to their respective maximum intensity along the profile. Corresponding local intensity maxima in both channels were used to assess recurrent local overlap of TcFabp8 and TcChs1 signals in subapical vesicle-like structures. Cuticle thickness was measured in Fiji (ImageJ). For each individual, three independent measurements were taken and averaged to obtain one biological replicate per elytron. Prior to statistical analysis, values were log2-transformed to stabilize variance. Differences between RNAi conditions were assessed using a Kruskal–Wallis test followed by pairwise Wilcoxon rank-sum tests with Benjamini-Hochberg correction for multiple comparisons. To account for potential non-independence of measurements within individuals, raw values were additionally analyzed using a linear mixed-effects model with treatment as fixed effect and individual as random intercept. This analysis yielded results consistent with those obtained from the aggregated data.

### 5.9. Expression and purification of recombinant TcFabp8

The *TcFabp8* variant 2 (*TcFabp8X2*) coding sequence was amplified from P4 pupal cDNA by PCR using extended primers to introduce *Nde*I and *BamHI* restriction sites for ligation into pET-16b. *E. coli* BL21 (pLysS) were transformed with the construct, and the resulting plasmid was purified and verified by DNA sequencing (Microsynth Seqlab).Expression was induced with 1 mM IPTG at OD_600_ ≈ 0.65 and continued for 3 h at 37 °C. Cells were harvested by centrifugation (4500 × g, 15 min, 4 °C), lysed by freeze–thawing and sonication (80 kHz, 40 % power, pulses), and the protein was purified from the soluble fraction using a HisTrap HP affinity column (Cytiva, USA) on an ÄKTA FPLC system (Amersham Biosystems, USA) with a 20–500 mM imidazole linear gradient. Elution fractions containing recombinant TcFabp8X2 were pooled, concentrated, and buffer-exchanged into PBS containing protease inhibitors. Success of expression and purification was controlled by SDS-PAGE and Western blotting.

### 5.10. X-ray diffraction (XRD) analysis of elytra after RNA interference

Fresh adult elytra were dissected after RNAi to silence *TcFABP8* or the *vermilion* gene *TcVER* as a control. For each measurement, one elytron from pupae injected with ds*TcVER* or ds*TcFABP8* were mounted side-by-side on silicon nitride supports and covered with 12.5 µm Kapton® foil to ensure identical exposure conditions. Three biological replicates per treatment were analyzed.

XRD data were acquired at the µSpot beamline (BESSY II, Berlin, Germany) using an 18 keV beam (λ = 0.689 Å) focused to ∼100 µm. Diffraction patterns were recorded in transmission geometry with a Dectris Eiger 9M detector at a sample-to-detector distance of 340 mm, covering a q-range of 0.10–41.78 nm⁻¹. Acquisition time was 10 s per position. To minimize beam damage, step sizes exceeded the beam footprint. Dioptas v0.7.1 [53] was used for detector calibration, mask generation, and visual inspection of diffraction patterns. Radial integration and background correction were performed using a pyFAI-based workflow with identical integration parameters for all samples [54]. For each diffraction pattern, the Kapton background was scaled individually using the Kapton-dominated region between the low-q fiber-correlation peak and the chitin 020 reflection. The scaling factor was estimated from q = 3.80 - 4.30 nm⁻¹ and constrained to avoid extended oversubtraction in the corrected profiles, while excluding the chitin peak regions from this quality-control criterion. Corrected one-dimensional intensity profiles, I(q), were then used for peak fitting and comparison of chitin-associated scattering features. Peak fitting was performed separately for the low-q correlation peak and the wider WAXS region. The low-q region was fitted from q = 1.10–1.55 nm⁻¹ using a power-law background and a pseudo-Voigt component for the fiber-correlation peak. The WAXS region was fitted from q = 4–26 nm⁻¹ using a linear background, broad Gaussian components to account for slowly varying background contributions, and pseudo-Voigt components for the major chitin-associated scattering features. All fits were performed identically for paired ds*TcVer* and ds*TcFABP8* samples, and fit-derived parameters were interpreted conservatively because some WAXS components represent overlapping peak envelopes rather than isolated Bragg reflections.

### 5.11. Azimuthal analysis of the low-q correlation signal

Local azimuthal anisotropy of the low-q correlation signal was quantified from background-corrected two-dimensional diffraction patterns using custom Python scripts [55]. The analysis followed the general principle of extracting local azimuthal intensity distributions from scanning X-ray diffraction patterns and representing their twofold directional component by a cosine-based model [56]. For each scan position, the Kapton background was subtracted after individual scale-factor correction as described above. Azimuthal intensity profiles, I(φ), were extracted in the q-range 1.30– 1.40 nm⁻¹, centered on the low-q correlation peak at q ≈ 1.36 nm⁻¹. Profiles were analyzed in detector-plane coordinates without 180° folding. This avoids artificial changes in apparent peak height or width that may arise when opposing lobes are not exactly antipodal or differ in shape due to projection geometry [57]. Following established approaches for describing twofold azimuthal orientation distributions by cosine functions [56], each local azimuthal profile was fitted with a second-order Fourier model:

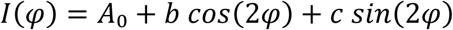

Here, I(φ) denotes the scattered intensity as a function of the azimuthal angle φ on the detector plane. The parameter A₀ represents the isotropic (angle-independent) background intensity component. The coefficients b and c describe the amplitude and phase of the anisotropic contribution with twofold symmetry (2φ), capturing directional variations in the scattering signal. The anisotropic amplitude was calculated as:

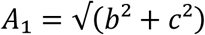

where A₁ represents the magnitude of the anisotropic component derived from the Fourier coefficients b and c. The degree of local detector-plane azimuthal anisotropy was defined as:

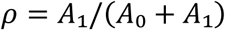

In this expression, ρ is a dimensionless parameter ranging between 0 and 1, where higher values indicate stronger anisotropy (i.e., more pronounced directional dependence of the scattering signal), and lower values indicate more isotropic scattering. The preferred detector-plane orientation of the 2φ component was calculated from the fitted Fourier coefficients and treated modulo 180° because of nematic symmetry. The resulting ρ values provide a dimensionless measure of local detector-plane azimuthal anisotropy, or projected local co-alignment strength, of the low-q correlation signal. They were not interpreted as an absolute three-dimensional fibril order parameter. Spatial maps of ρ and the corresponding detector-plane orientation vectors were reconstructed from the original scan geometry and analyzed separately for each paired scan and treatment [56].

### 5.12. Statistical Analysis

All quantitative data were obtained from at least three biological replicates. Values are presented as mean ± standard deviation (SD) unless otherwise indicated. Statistical analyses were performed in R (version 4.2.2). Depending on data distribution and variance homogeneity, either parametric or non-parametric tests were applied. For comparisons involving multiple groups, differences were assessed using the Kruskal-Wallis test followed by pairwise Wilcoxon rank-sum tests with Benjamini-Hochberg correction for multiple comparisons. Where appropriate, linear mixed-effects models were used to account for non-independence of measurements within individuals. Significance was set at p < 0.05.

## 6. Declarations

### 6.1. Ethics approval and consent to participate

Not applicable

### 6.2. Consent for publication

Not applicable

### 6.3. Availability of data and materials

The datasets used and/or analysed during the current study are available from the corresponding author on reasonable request. Custom analysis scripts are available in the Zenodo repository: https://doi.org/10.5281/zenodo.21525203

### 6.4. Competing interests

The authors declare that they have no competing interests.

### 6.5. Funding

This work was supported by DFG grants ME2210/9-1 received in the scope of Joint Sino-German Research Projects 2021 as well as ME2210/11-1 of the DFG priority programme 2416 “Codeχ”.

### 6.6. Authors’ contributions

CRediT authorship contribution statement

**MB:** Conceptualization, Data curation, Formal analysis, Investigation, Methodology, Software, Supervision, Validation, Visualization, Writing – original draft, Writing – review & editing.

**KV:** Data curation, Investigation, Visualization.

**WZ:** Methodology, Validation, Writing – review & editing.

**LB:** Formal analysis, Software, Validation.

**YP:** Formal analysis, Validation, Writing – review & editing.

**HM:** Conceptualization, Formal analysis, Funding acquisition, Methodology, Project administration, Supervision, Validation, Writing – review & editing. All authors read and approved the final manuscript.

### 6.7. Acknowledgements

We thank Carolin Fisher and Ernesto Scopolla for their help with X-ray diffraction analysis. We further acknowledge the Helmholtz-Zentrum Berlin (HZB) for the allocation of synchrotron radiation beamtime and hosting. X-ray diffraction data were collected at the mySpot beamline operated at the BESSY II synchrotron (Berlin- Adlershof, Germany).

## Notes

### Competing Interest Statement

The authors have declared no competing interest.

