## Supplemental Information for "A fatty acid-binding protein links lipid handling to chitin synthase-dependent cuticle formation and nanoscale organization"

#### SI Appendix: Results

**Figure S1** AlphaFold model TcFabp8X1, TcFabp8X2 and TcFabp8X3

(A) Predicted structures of TcFabp8X1, TcFabp8X2, and TcFabp8X3 showing the conserved FABP β-barrel fold. TcFABP8X1 contains an N-terminal Sec/SPI signal peptide (dark green), TcFbap8X2 represents the core cytosolic form, and TcFabp8X3 exhibits an alternative N-terminal sequence (magenta). (B) pLDDT values aligned to Fabp core domain

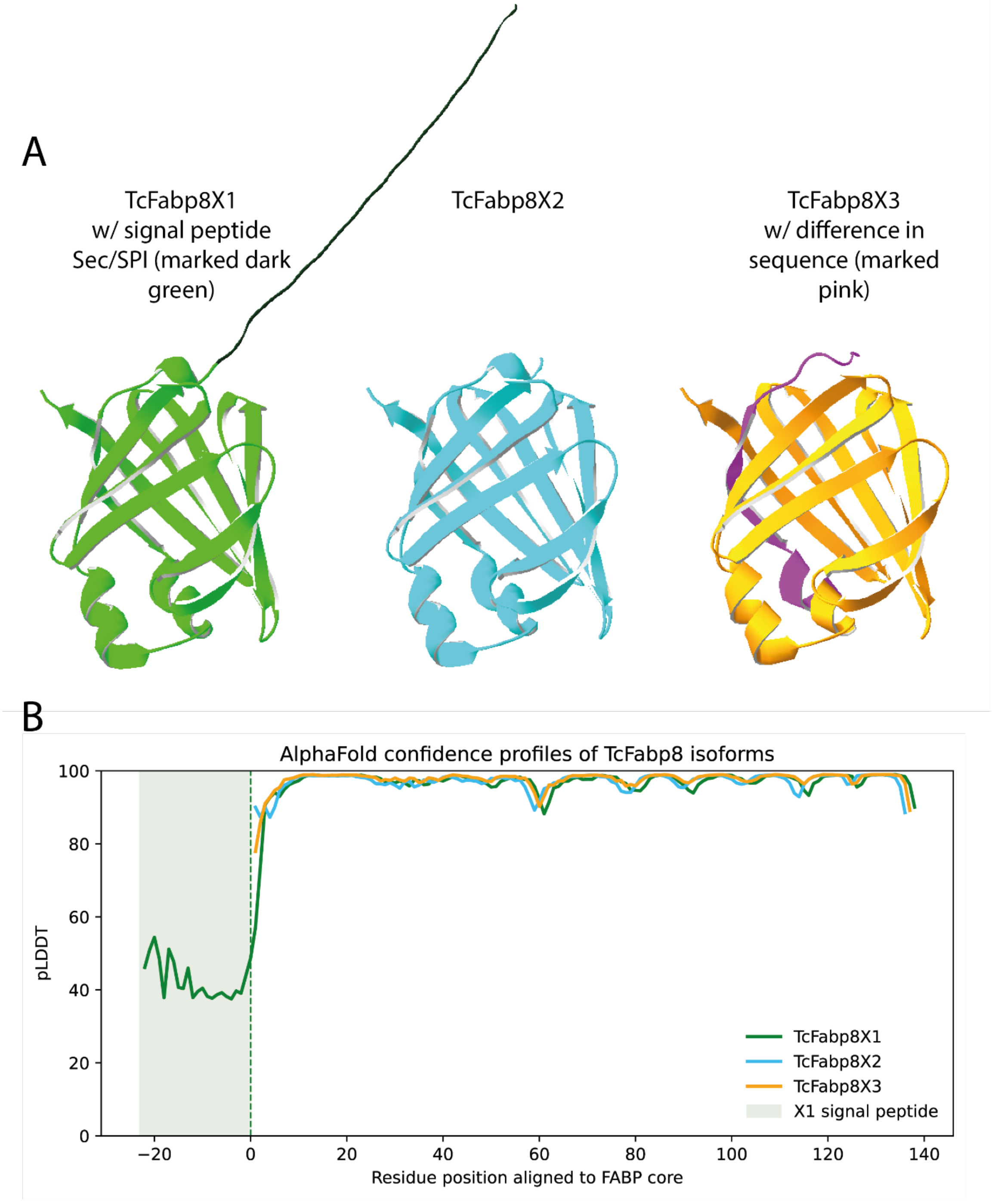

**Figure S2** MSA (MAFFT) basis for phylogenetics

Multiple sequence alignment generated with MAFFT of TcFabp8 isoforms and representative Fabp homologs used for phylogenetic analysis. Conserved residues within the Fabp core domain are highlighted, whereas most sequence variation is confined to the N-terminal region, including the signal peptide of TcFabp8X1.

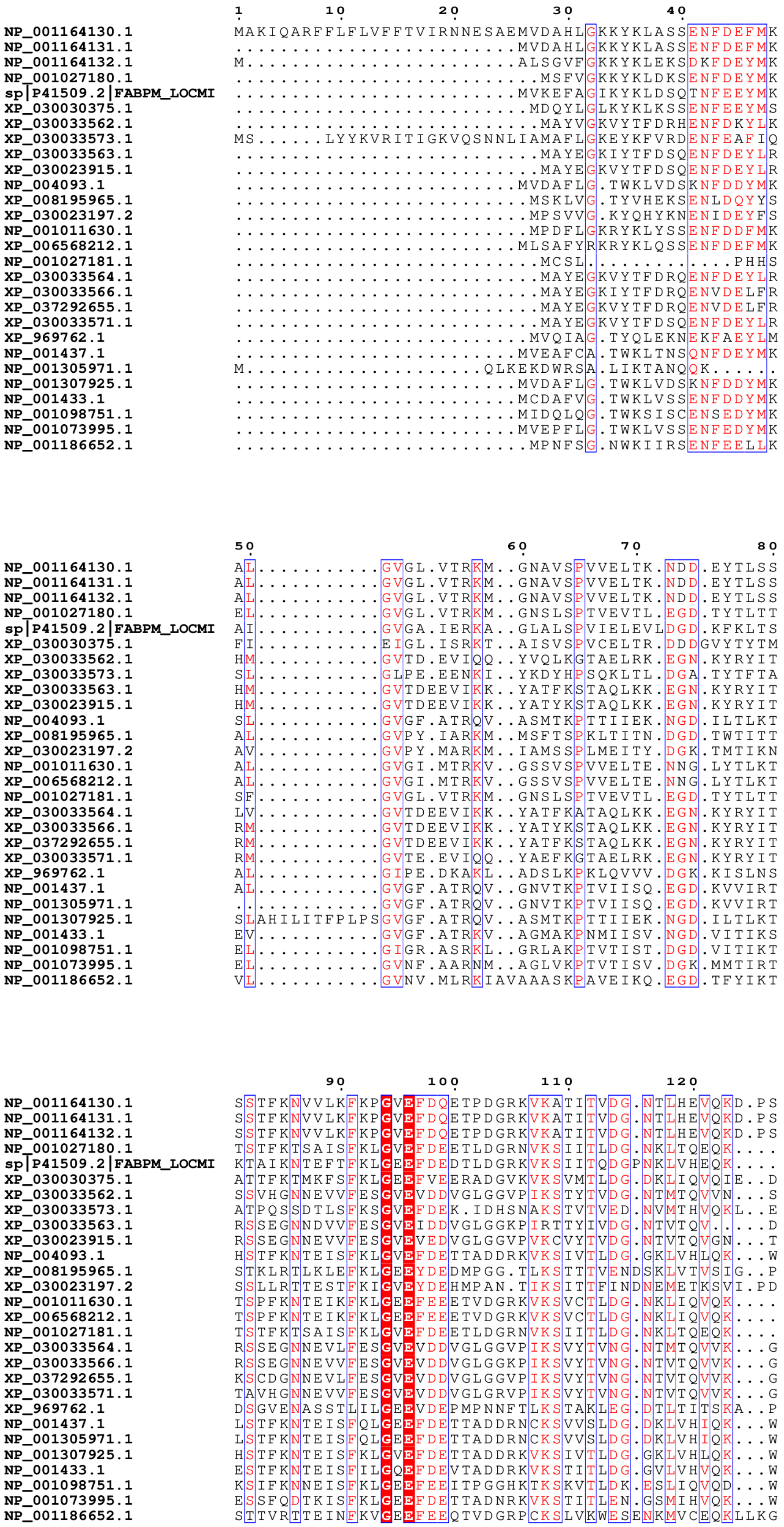
1 10 20 30 40
NP_001164130.1 MAKIQARFFLFLVFFTVIRNNESAEMVDAHLGKKYKLASSENFDEFMK
NP_001164131.1 .........................MVDAHLGKKYKLASSENFDEFMK
NP_001164132.1 M........................ALSGVFGKKYKLEKSDKFDEYMK
NP_001027180.1 ...........................MSFVGKKYKLDKSENFDEYMK
sp|P41509.2|FABPM_LOCMI .........................MVKEFAGIKYKLDSQTNFEEYMK
XP_030030375.1 ..........................MDQYLGLKYKLKSSENFEEYMS
XP_030033562.1 ...........................MAYVGKVYTFDRHENFDKYLK
XP_030033573.1 MS......LYYKVRITIGKVQSNNLIAMAFLGKEYKFVRDENFEAFIQ
XP_030033563.1 ...........................MAYEGKIYTFDSQENFDEYLR
XP_030023915.1 ...........................MAYEGKVYTFDSQENFDEYLR
NP_004093.1 .........................MVDAFLG.TWKLVDSKNFDDYMK
XP_008195965.1 ..........................MSKLVG.TYVHEKSENLDQYYS
XP_030023197.2 ..........................MPSVVG.KYQHYKNENIDEYFS
NP_001011630.1 ..........................MPDFLGKRYKLYSSENFDDFMK
XP_006568212.1 .........................MLSAFYRKRYKLQSSENFDEFMK
NP_001027181.1 ...........................MCSL.............PHHS
XP_030033564.1 ...........................MAYEGKVYTFDRQENFDEYLR
XP_030033566.1 ...........................MAYEGKIYTFDRQENVDELFR
XP_037292655.1 ...........................MAYEGKVYTFDRQENVDELFR
XP_030033571.1 ...........................MAYEGKVYTFDSQENFDEYLR
XP_969762.1 ..........................MVQIAG.TYQLEKNEKFAEYLM
NP_001437.1 .........................MVEAFCA.TWKLTNSQNFDEYMK
NP_001305971.1 M.....................QLKEKDWRSA.LIKTANQQK......
NP_001307925.1 .........................MVDAFLG.TWKLVDSKNFDDYMK
NP_001433.1 .........................MCDAFVG.TWKLVSSENFDDYMK
NP_001098751.1 .........................MIDQLQG.TWKSISCENSEDYMK
NP_001073995.1 .........................MVEPFLG.TWKLVSSENFEDYMK
NP_001186652.1 ..........................MPNFSG.NWKIIRSENFEELLK
50 60 70 80
NP_001164130.1 AL...........GVGL.VTRKM..GNAVSPVVELTK.NDD.EYTLSS
NP_001164131.1 AL...........GVGL.VTRKM..GNAVSPVVELTK.NDD.EYTLSS
NP_001164132.1 AL...........GVGL.VTRKM..GNAVSPVVELTK.NDD.EYTLSS
NP_001027180.1 EL...........GVGL.VTRKM..GNSLSPTVEVTL.EGD.TYTLTT
sp|P41509.2|FABPM_LOCMI AI...........GVGA.IERKA..GLALSPVIELEVLDGD.KFKLTS
XP_030030375.1 FI...........EIGL.ISRKT..AISVSPVCELTR.DDDGVYTYTM
XP_030033562.1 HM...........GVTD.EVIQQ..YVQLKGTAELRK.EGN.KYRYIT
XP_030033573.1 SL...........GLPE.EENKI..YKDYHPSQKLTL.DGA.TYTFTA
XP_030033563.1 HM...........GVTDEEVIKK..YATFKSTAQLKK.EGN.KYRYIT
XP_030023915.1 HM...........GVTDEEVIKK..YATYKSTAQLKK.EGN.KYRYIT
NP_004093.1 SL...........GVGF.ATRQV..ASMTKPTTIIEK.NGD.ILTLKT
XP_008195965.1 AL...........GVPY.IARKM..MSFTSPKLTITN.DGD.TWTITT
XP_030023197.2 AV...........GVPY.MARKM..IAMSSPLMEITY.DGK.TMTIKN
NP_001011630.1 AL...........GVGI.MTRKV..GSSVSPVVELTE.NNG.LYTLKT
XP_006568212.1 AL...........GVGI.MTRKV..GSSVSPVVELTE.NNG.LYTLKT
NP_001027181.1 SF...........GVGL.VTRKM..GNSLSPTVEVTL.EGD.TYTLTT
XP_030033564.1 LV...........GVTDEEVIKK..YATFKATAQLKK.EGN.KYRYIT
XP_030033566.1 RM...........GVTDEEVIKK..YATYKSTAQLKK.EGN.KYRYIT
XP_037292655.1 RM...........GVTDEEVIKK..YATFKSTAQLKK.EGN.KYRYIT
XP_030033571.1 RM...........GVTE.EVIQQ..YAEFKGTAELRK.EGN.KYRYIT
XP_969762.1 AL...........GIPE.DKAKL..ADSLKPKLQVVV.DGK.KISLNS
NP_001437.1 AL...........GVGF.ATRQV..GNVTKPTVIISQ.EGD.KVVIRT
NP_001305971.1 .............GVGF.ATRQV..GNVTKPTVIISQ.EGD.KVVIRT
NP_001307925.1 SLAHILITFPLPSGVGF.ATRQV..ASMTKPTTIIEK.NGD.ILTLKT
NP_001433.1 EV...........GVGF.ATRKV..AGMAKPNMIISV.NGD.VITIKS
NP_001098751.1 EL...........GIGR.ASRKL..GRLAKPTVTIST.DGD.VITIKT
NP_001073995.1 EL...........GVNF.AARNM..AGLVKPTVTISV.DGK.MMTIRT
NP_001186652.1 VL...........GVNV.MLRKIAVAAASKPAVEIKQ.EGD.TFYIKT
90 100 110 120
NP_001164130.1 SSTFKNVVLKFKPGVEFDQETPDGRKVKATITVDG.NTLHEVQKD.PS
NP_001164131.1 SSTFKNVVLKFKPGVEFDQETPDGRKVKATITVDG.NTLHEVQKD.PS
NP_001164132.1 SSTFKNVVLKFKPGVEFDQETPDGRKVKATITVDG.NTLHEVQKD.PS
NP_001027180.1 TSTFKTSAISFKLGVEFDEETLDGRNVKSIITLDG.NKLTQEQK....
sp|P41509.2|FABPM_LOCMI KTAIKNTEFTFKLGEEFDEDTLDGRKVKSIITQDGPNKLVHEQK....
XP_030030375.1 ATTFKTMKFSFKLGEEFVEERADGVKVKSVMTLDG.DKLIQVQIE..D
XP_030033562.1 SSVHGNNEVVFESGVEVDDVGLGGVPIKSTYTVDG.NTMTQVVN...S
XP_030033573.1 ATPQSSDTLSFKSGVEFDEK.IDHSNAKSTVTVED.NVMTHVQKL..E
XP_030033563.1 RSSEGNNDVVFESGVEIDDVGLGGKPIRTTYIVDG.NTVTQV.....D
XP_030023915.1 RSSEGNNEVVFESGVEVEDVGLGGVPVKCVYTVDG.NTVTQVGN...T
NP_004093.1 HSTFKNTEISFKLGVEFDETTADDRKVKSIVTLDG.GKLVHLQK...W
XP_008195965.1 STKLRTLKLEFKLGEEYDEDMPGG.TLKSTTTVENDSKLVTVSIG..P
XP_030023197.2 SSLLRTTESTFKIGVEYDEHMPAN.TIKSITFINDNEMETKSVI.PD
NP_001011630.1 TSPFKNTEIKFKLGEEFEEETVDGRKVKSVCTLDG.NKLIQVQK....
XP_006568212.1 TSPFKNTEIKFKLGEEFEEETVDGRKVKSVCTLDG.NKLIQVQK....
NP_001027181.1 TSTFKTSAISFKLGVEFDEETLDGRNVKSIITLDG.NKLTQEQK....
XP_030033564.1 RSSEGNNEVLFESGVEVDDVGLGGVPIKSVYTVNG.NTMTQVVK...G
XP_030033566.1 RSSEGNNEVVFESGVEVDDVGLGGKPIKSVYTVNG.NTVTQVVK...G
XP_037292655.1 KSCDGNNEVLFESGVEVDDVGLGGVPIKSVYTVNG.NTVTQVVK...G
XP_030033571.1 TAVHGNNEVVFESGVEVDDVGLGRVPIKSVYTVNG.NTVTQVVK...G
XP_969762.1 DSGVENASSTLILGEEVDEPMPNNFTLKSTAKLEG.DTLTITSKA..P
NP_001437.1 LSTFKNTEISFQLGEEFDETTADDRNCKSVVSLDG.DKLVHIQK...W
NP_001305971.1 LSTFKNTEISFQLGEEFDETTADDRNCKSVVSLDG.DKLVHIQK...W
NP_001307925.1 HSTFKNTEISFKLGVEFDETTADDRKVKSIVTLDG.GKLVHLQK...W
NP_001433.1 ESTFKNTEISFILGQEFDEVTADDRKVKSTITLDG.GVLVHVQK...W
NP_001098751.1 KSIFKNNEISFKLGEEFEEITPGGHKTKSKVTLDK.ESLIQVQD...W
NP_001073995.1 ESSFQDTKISFKLGEEFDETTADNRKVKSTITLEN.GSMIHVQK...W
NP_001186652.1 STTVRTTEINFKVGEEFEEQTVDGRPCKSLVKWESENKMVCEQKLLKG

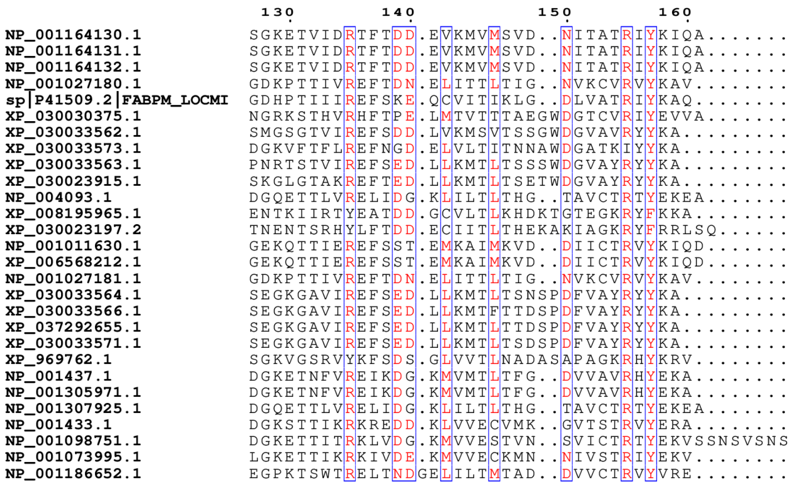
130 140 150 160
NP_001164130.1 SGKETVIDRTFTDD.EVKMVMSVD..NITATRIYKIQA.......
NP_001164131.1 SGKETVIDRTFTDD.EVKMVMSVD..NITATRIYKIQA.......
NP_001164132.1 SGKETVIDRTFTDD.EVKMVMSVD..NITATRIYKIQA.......
NP_001027180.1 GDKPTTIVREFTDN.ELITTLTIG..NVKCVRVYKAV........
sp|P41509.2|FABPM_LOCMI GDHPTIIIREFSKE.QCVITIKLG..DLVATRIYKAQ........
XP_030030375.1 NGRKSTHVRHFTPE.LMTVTTTAEGWDGTCVRIYEVVA.......
XP_030033562.1 SMGSGTVIREFSDD.LVKMSVTSSGWDGVAVRYYKA.........
XP_030033573.1 DGKVFTFLREFNGD.ELVLTITNNAWDGATKIYYKA.........
XP_030033563.1 PNRTSTVIREFSED.LLKMTLTSSSWDGVAYRYYKA.........
XP_030023915.1 SKGLGTAKREFTED.LLKMTLTSETWDGVAYRYYKA.........
NP_004093.1 DGQETTLVRELIDG.KLILTLTHG..TAVCTRTYEKEA.......
XP_008195965.1 ENTKIIRTYEATDD.GCVLTLKHDKTGTEGKRYFKKA........
XP_030023197.2 TNENTSRHYLFTDD.ECIITLTHEKAKIAGKRYFRRLSQ......
NP_001011630.1 GEKQTTIEREFSST.EMKAIMKVD..DIICTRVYKIQD.......
XP_006568212.1 GEKQTTIEREFSST.EMKAIMKVD..DIICTRVYKIQD.......
NP_001027181.1 GDKPTTIVREFTDN.ELITTLTIG..NVKCVRVYKAV........
XP_030033564.1 SEGKGAVIREFSED.LLKMTLTSNSPDFVAYRYYKA.........
XP_030033566.1 SEGKGAVIREFSED.LLKMTFTTDSPDFVAYRYYKA.........
XP_037292655.1 SEGKGAVIREFSED.LLKMTLTTDSPDFVAYRYYKA.........
XP_030033571.1 SEGKGAVIREFSED.LLKMTLTSDSPDFVAYRYYKA.........
XP_969762.1 SGKVGSRVYKFSDS.GLVVTLNADASAPAGKRHYKRV........
NP_001437.1 DGKETNFVREIKDG.KMVMTLTFG..DVVAVRHYEKA........
NP_001305971.1 DGKETNFVREIKDG.KMVMTLTFG..DVVAVRHYEKA........
NP_001307925.1 DGQETTLVRELIDG.KLILTLTHG..TAVCTRTYEKEA.......
NP_001433.1 DGKSTTIKRKREDD.KLVVECVMK..GVTSTRVYERA........
NP_001098751.1 DGKETTITRKLVDG.KMVVESTVN..SVICTRTYEKVSSNSVSNS
NP_001073995.1 LGKETTIKRKIVDE.KMVVECKMN..NIVSTRIYEKV........
NP_001186652.1 EGPKTSWTRELTNDGELILTMTAD..DVVCTRVYVRE........

**Figure S3** SignalP predictions for TcFabp8X1, TcFabp8X2 and TcFabp8X3 variants

SignalP 6.0 analysis of TcFabp8X1, TcFabp8X2, and TcFabp8X3. TcFabp8X1 shows a predicted Sec/SPI signal peptide with a defined cleavage site, whereas TcFabp8X2 and TcFabp8X3 lack signal peptide features and are predicted as non-secretory proteins.

### TcFabp8X1

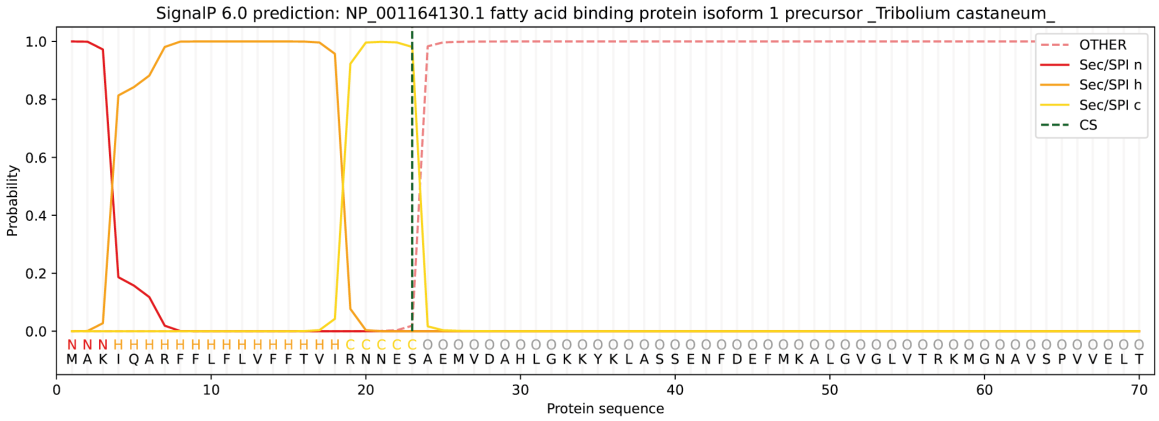


### TcFabp8X2

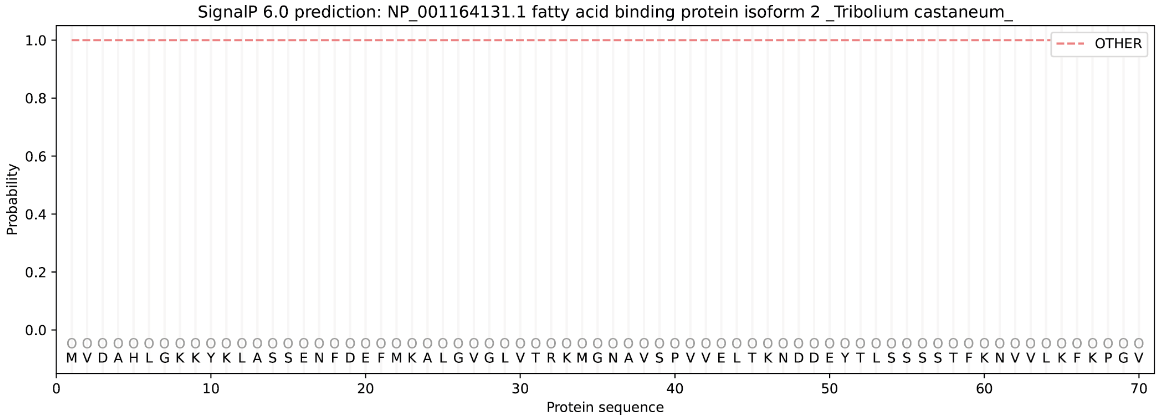


### TcFabp8X3

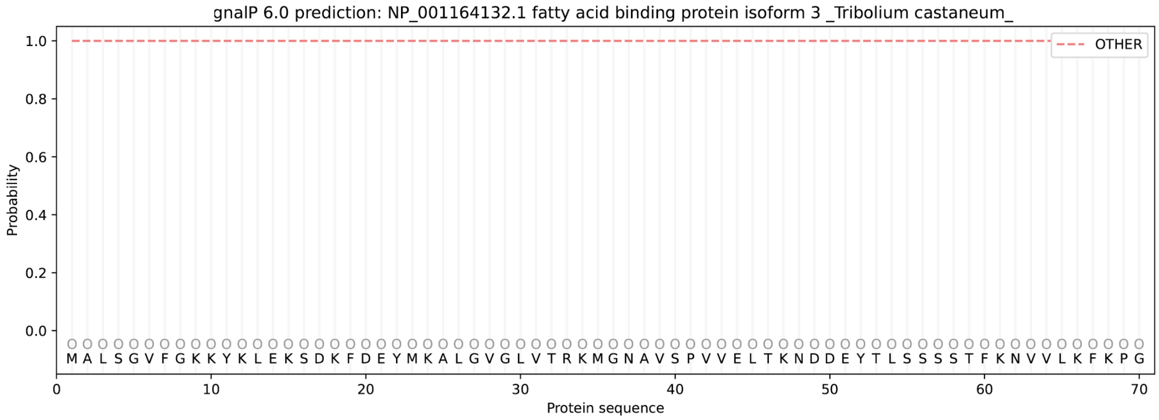

**Figure S4** Expression and purification of recombinant TcFabp8X2.

SDS-PAGE analysis of recombinant TcFabp8X2 showing expression and enrichment of the protein at the expected molecular weight of ~ 16 kDa.

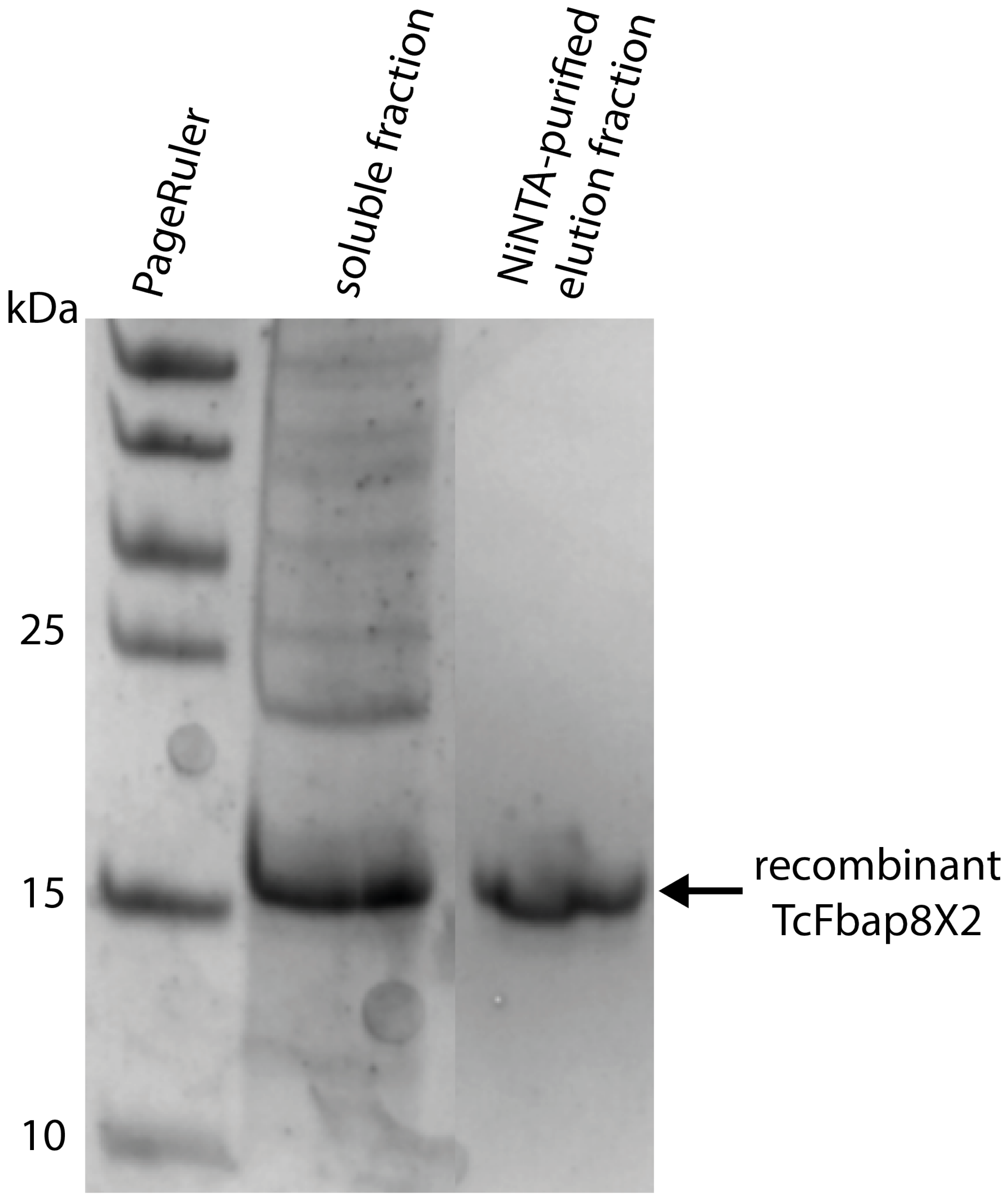

**Figure S5** Developmental expression of *TcFABP8* splice variants and *TcCHS1*.

qPCR analysis of *TcCHS1* and *TcFABP8* isoform expression across pupal stages P0–P4 (Fig. 2A). Expression is shown as log2(normalized Ct). *TcFABP8X2* and *TcCHS1* display coordinated upregulation toward P4, whereas *TcFABP8X3* remains low and uncorrelated.

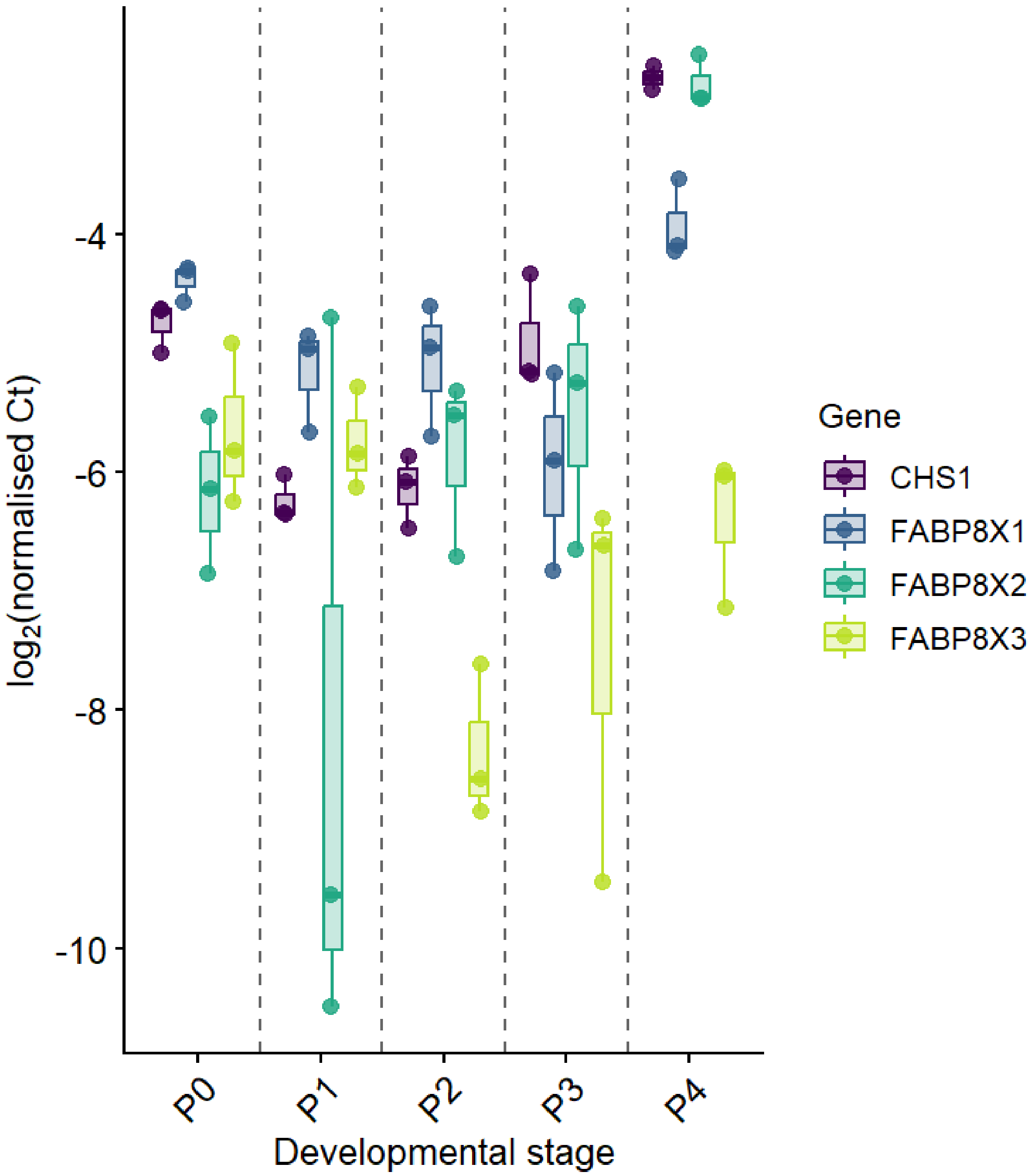

**Figure S6** RNAi efficiencies of *TcCHS1, TcFABP2, TcFABP8* and *TcTH*

qPCR-based assessment of knockdown efficiencies for ds*TcCHS1*, ds*TcFABP2*, ds*TcFABP8*, and ds*TcTH*. Relative expression levels are shown as percentage of control, demonstrating efficient depletion of target transcripts in all RNAi treatments.

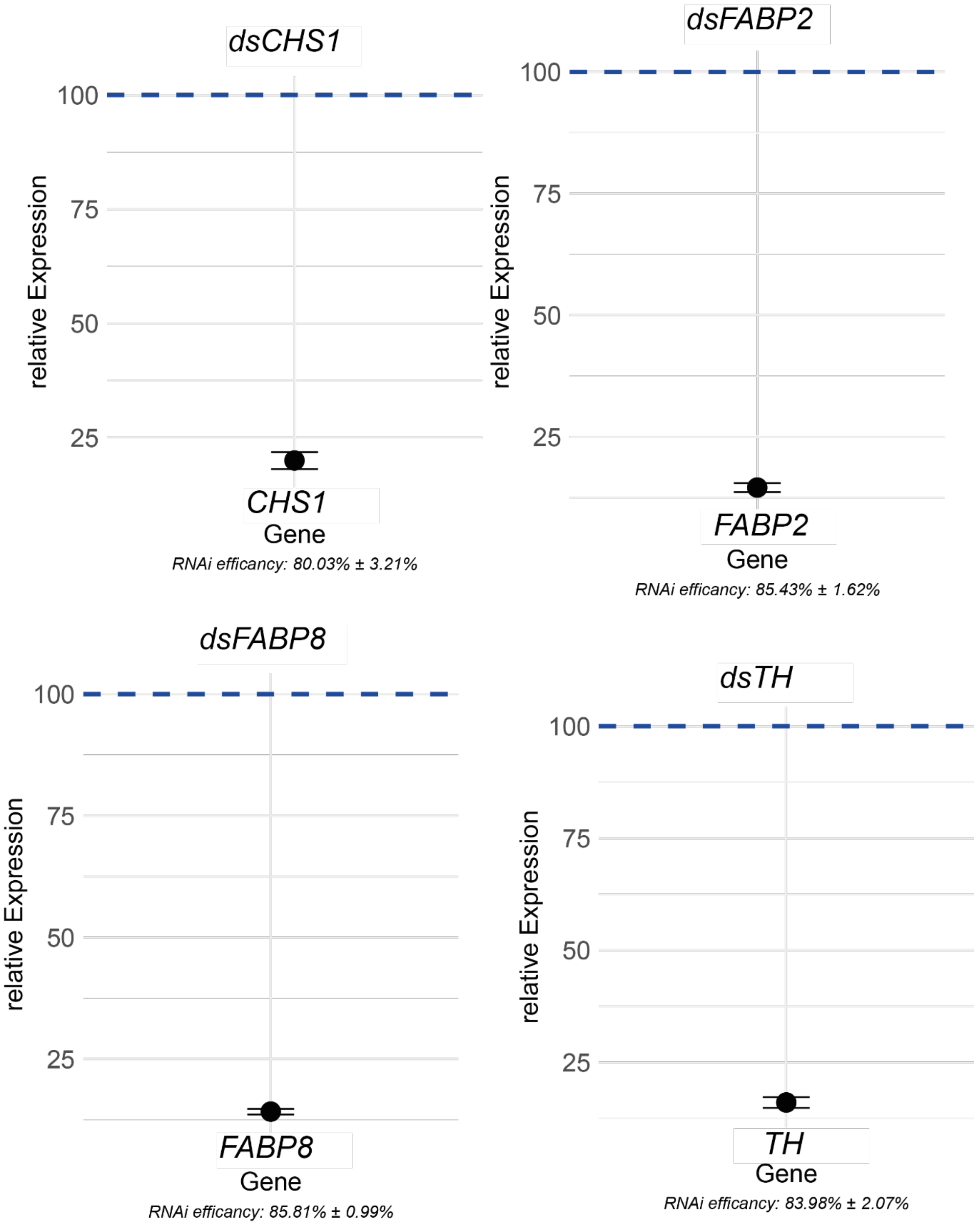

**Figure S7** Representative peak fitting of one-dimensional X-ray diffraction profiles from elytral cuticle.

Representative background-corrected XRD intensity profiles are shown for three biological replicate pairs of ds*TcFABP8*- and ds*TcVER*-treated adult elytra. Orange points indicate experimental data, and the green line shows the combined fit. Individual fit components are shown separately, including Gaussian background/low-frequency contributions, a linear baseline, and pseudo-Voigt functions used to model the major α-chitin-associated reflections in the q-range of approximately 3–26 nm$^{-1}$. The fitted components reproduce the characteristic chitin reflections, including the broad peak around q ≈ 14 nm$^{-1}$ and sharper high-q reflections around q ≈ 18–19 nm$^{-1}$. Rows correspond to biological replicates; columns show the paired RNAi conditions measured under comparable acquisition and background-correction settings. Intensity is shown in arbitrary units.

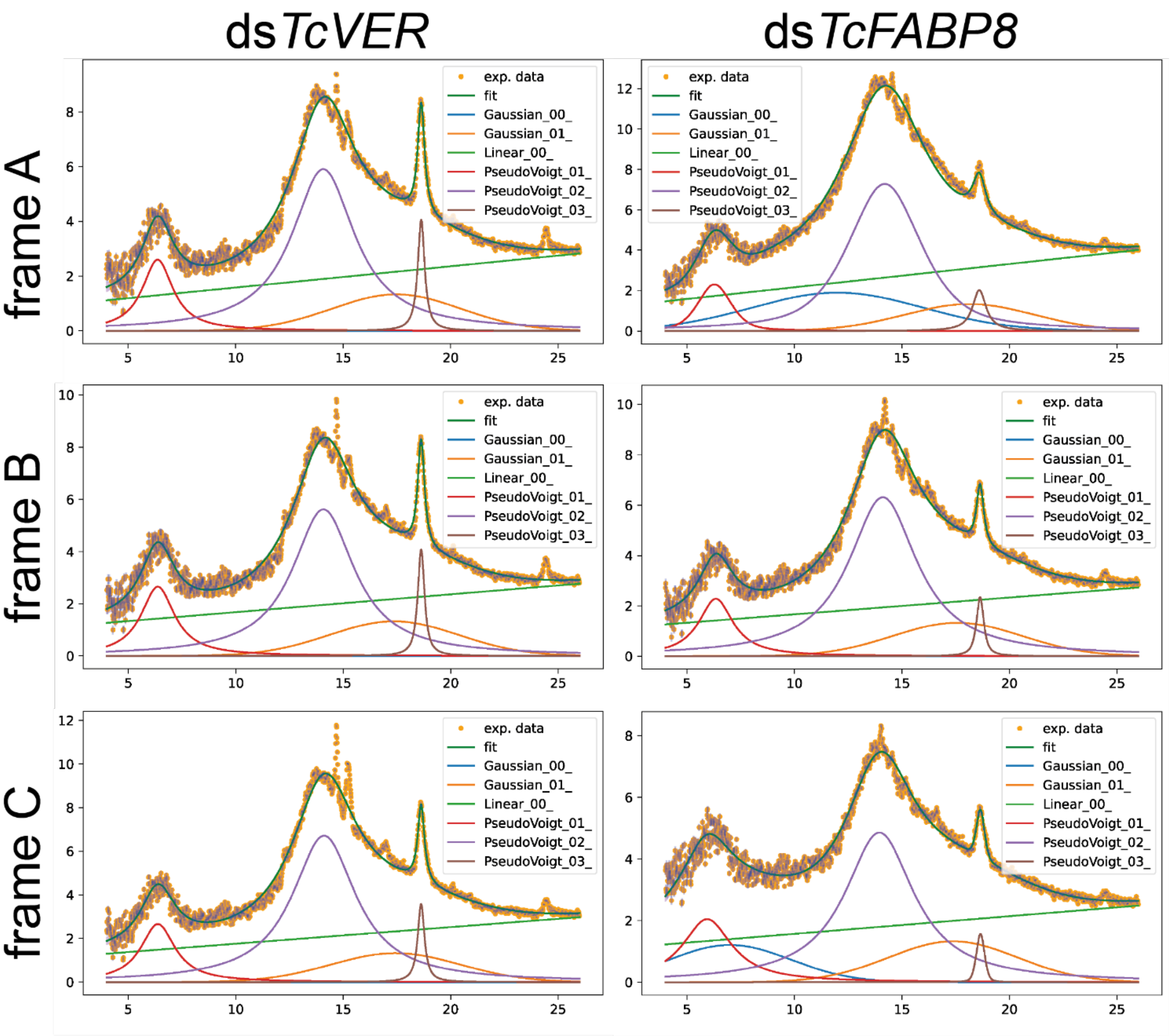

Supplement tables

**Table S1** Oligos used in this study

| **Name** | **Sequence 5' -> 3'** |
|---|---|
| **T7_*FABP*8_fwd** | TAATACGACTCACTATAGGGCGACGTTCAAAAACGTGGT |
| **T7_*FABP*8_rev** | TAATACGACTCACTATAGGGTACGGGTGGCTGTGATGTTA |
| **q*FABP*8X3_fwd** | AACCCGAACTTTAGGCTGTGA |
| **q*FABP*8_rev** | TTTGAACGTCGAGGACGAACT |
| **T7_*FABP*2_fwd** | TAATACGACTCACTATAGGGTGTCGAAACTTGTGGGAACA |
| **T7_*FABP*2_rev** | TAATACGACTCACTATAGGGGGCATGTCCTCGTCGTATTC |
| **q*FABP*2_fwd** | ACTGTATCTTGCAGTAGGTTGAC |
| **q*FABP*2_rev** | AGTCCAAGTATCGCCGTCATT |
| **T7_*FABP*4_fwd** | TAATACGACTCACTATAGGGTTGGAATAGCACAGCACAGC |
| **T7_*FABP*4_rev** | TAATACGACTCACTATAGGGGCGTTCAAAGTCACAACGAG |
| **q*FABP*4_fwd** | GGCGAGGAAGTAGACGAACC |
| **q*FABP*4_rev** | TGTAGTGCCTCTTTCCAGCG |
| **RpS18_fwd** | CGAAGAGGTCGAGAAAATCG |
| **RpS18_rev** | CGTGGTCTTGGTGTGTTGAC |
| **qCHS1_fwd** | GTCTTCTTCACCGAAACGCTCC |
| **qCHS1_rev** | CGCGACTTCTCCTTCTTTGACC |
| **T7_VER_fwd** | TAATACGACTCACTATAGGTTGGTGGACCACGAAATGAT |

| | |
|---|---|
| **T7_VER_rev** | TAATACGACTCACTATAGGGCCATTTCGTAATCAGCGAG |
| ***FABP*8_FL_NdeI_fwd** | TTTTCATATGGTCGACGCACATCTTGGCAAG |
| ***FABP*8_FL_BamHI_rev** | TTTTGGATCCTTAGGCTTGAATCTTGTAAATACGGGTGG |
| **q*FABP*8_X1_fwd** | CCAAAATCCAAGCGCGATTCT |
| **q*FABP*8_X2_fwd** | TCGCGTAATAGACGCCTTCG |
| **T7-CHS1-fwd** | TAATACGACTCACTATAGGGAATCGATTTGTTCGCAATTG |
| **T7-CHS1-rev** | TAATACGACTCACTATAGGGCGAAAATGTAGGCGAAATA |
| **T7-TH-fwd** | TAATACGACTCACTATAGGGGTGGCTGCTGCTCAAAAAAATC |
| **T7-TH-rev** | TAATACGACTCACTATAGGGTCAGGCCAGAATCGTCTTCG |
| **qTH_fwd** | AGTGAGATTCAACCCGCACACC |
| **qTH_rev** | ATGAGCCGATGAAAGCACGAGG |
| **qLAC2_fwd** | ATTGGATCCAACTTCGAGGC |
| **qLAC2_rev** | GCACATGCAGTTTTGTTGGC |

**Table S2** Antibodies used in this study

| Antibody | Host | Source | Dilution |
|---|---|---|---|
| **αChs1-C** | Rabbit | Immunized against peptide by Proteogenix | 1:50 |
| **αChs1-ATTO647N** | Rabbit | αChs1-C coupled to ATTO647N | 1:50 |
| **αRabbit-Cy3** | Goat | Commercial Cy3-conjugated antibody (Thermo) | 1:1000 |
| **αFABP4** | Rabbit | Commercial antibody agains human FABP4 (Proteogenix, France) | 1:50 |
| **αRabbit-FITC** | Goat | Commercial FITC-conjugated antibody (Thermo) | 1:500 |
| **αHIS** | Mouse | Commercial antibody (Thermo) | 1:6000 |
| **αMouse** | Goat | Commercial AP-conjugated antibody (Thermo) | 1:6000 |

**Table S3** Dyes used in this study

| Dye | Target | Source | Dilution |
|---|---|---|---|
| **CBD-FITC** | Chitin | CBD recombinant expressed in lab, FITC-maleimid by Thermo Fisher | 1:250 |
| **DAPI** | Nuclei | Thermo Fisher | 1:500 |
| **Phalloidin-ATTO633** | F-actin | ATTO-Tec, Siegen | 1:100 |
| **Nile Red** | Lipids | Thermo Fisher | 10µg/ml |
| **Calcofluor White** | Chitin | Thermo Fisher | 10 µg/ml |
| **ATTO647-NHS ester** | Antibody labeling | ATTO-Tec, Siegen | 10 fold molar excess |

**Table S4** SignalP6.0 predicition of signal peptides in TcFabp8 isoforms

| **# SignalP-6.0** | **Organism: Eukarya** | **Timestamp: 20260203102432** | | |
|---|---|---|---|---|
| **# ID** | Prediction | OTHER | SP(Sec/SPI) | CS Position |
| **TcFabp8X1 (NP_001164130.1)** | SP | 0.000151 | 0.999850 | CS pos: 23-24. Pr: 0.9810 |
| **TcFabp8X2 (NP_001164131.1)** | OTHER | 1.000000 | 0.000000 | |
| **TcFabp8X3 (NP_001164132.1)** | OTHER | 1.000000 | 0.000000 | |

Table S5 **Low-q correlation peak fit parameters after scaled Kapton-background subtraction.**

Low-q profiles were fitted from q = 1.10–1.55 $nm^{-1}$ using a power-law background and one pseudo-Voigt component for the low-q correlation peak. d-spacings were calculated as d = 2π/q. Values are shown for paired ds*TcVER*and ds*TcFABP8* elytra measured in the same frame.

| **Frame** | **Treatment** | **q center [$nm^{-1}$]** | **d = 2π/q [nm]** | **σ [$nm^{-1}$]** | **FWHM_q [$nm^{-1}$]** | **Amplitude [a.u.]** |
|---|---|---|---|---|---|---|
| A | ds*TcVER* | 1.3495 | 4.6560 | 0.0642 | 0.1511 | 2.982 |
| A | ds*TcFABP8* | 1.3303 | 4.7232 | 0.0763 | 0.1797 | 4.253 |
| B | ds*TcVER* | 1.3514 | 4.6493 | 0.0577 | 0.1360 | 3.410 |
| B | ds*TcFABP8* | 1.3367 | 4.7003 | 0.0620 | 0.1459 | 3.424 |
| C | ds*TcVER* | 1.3519 | 4.6476 | 0.0557 | 0.1313 | 4.722 |
| C | ds*TcFABP8* | 1.3055 | 4.8130 | 0.1344 | 0.3165 | 5.611 |

**Table S6** Summary of low-q correlation peak parameters. Values represent mean ± SD across the three paired frames.

| Parameter | ds*TcVER* mean ± SD | ds*TcFABP8* mean ± SD | Interpretation |
|---|---|---|---|
| q center $[nm^{-1}]$ | 1.3509 ± 0.0013 | 1.3242 ± 0.0165 | Lower q after TcFabp8 depletion |
| d = 2π/q [nm] | 4.6510 ± 0.0044 | 4.7454 ± 0.0594 | Larger correlation distance |
| σ $[nm^{-1}]$ | 0.0592 ± 0.0044 | 0.0909 ± 0.0384 | Broader low-q peak, strongest in frame C |
| FWHM_q $[nm^{-1}]$ | 0.1395 ± 0.0104 | 0.2140 ± 0.0903 | Broader low-q peak |
| Amplitude [a.u.] | 3.705 ± 0.906 | 4.429 ± 1.105 | Not consistently reduced |

**Table S7** WAXS pseudo-Voigt component fit summary after scaled Kapton-background subtraction.

WAXS profiles were fitted from q = 4–26 $nm^{-1}$. Components are interpreted conservatively because several fitted components represent broad or overlapping peak envelopes rather than isolated Bragg reflections. Values represent mean ± SD across three paired frames.

| Component | Approximate region | Parameter | ds*TcVER* mean ± SD | ds*TcFABP8* mean ± SD |
|---|---|---|---|---|
| PV01 | ~6.3 $nm^{-1}$ / 020-region | q center [$nm^{-1}$] | 6.3714 ± 0.0050 | 6.1861 ± 0.2183 |
| PV01 | ~6.3 $nm^{-1}$ / 020-region | σ [$nm^{-1}$] | 0.9290 ± 0.0551 | 1.0881 ± 0.2650 |
| PV01 | ~6.3 $nm^{-1}$ / 020-region | Amplitude [a.u.] | 7.704 ± 0.556 | 7.074 ± 1.816 |
| PV02 | ~14.1 $nm^{-1}$ / 110/120 envelope | q center [$nm^{-1}$] | 14.0850 ± 0.0194 | 14.0866 ± 0.1268 |
| PV02 | ~14.1 $nm^{-1}$ / 110/120 envelope | σ [$nm^{-1}$] | 1.7782 ± 0.0252 | 1.9702 ± 0.0370 |
| PV02 | ~14.1 $nm^{-1}$ / 110/120 envelope | Amplitude [a.u.] | 33.985 ± 3.601 | 36.117 ± 5.431 |
| PV03 | ~18.6 $nm^{-1}$ / 013-region | q center [$nm^{-1}$] | 18.6244 ± 0.0036 | 18.6265 ± 0.0273 |
| PV03 | ~18.6 $nm^{-1}$ / 013-region | σ [$nm^{-1}$] | 0.1894 ± 0.0019 | 0.2671 ± 0.0950 |
| PV03 | ~18.6 $nm^{-1}$ / 013-region | Amplitude [a.u.] | 2.303 ± 0.170 | 1.555 ± 0.777 |

**Table S8** Summary of local detector-plane azimuthal anisotropy parameters for the low-q correlation signal in paired ds*TcVER* and ds*TcFABP8* elytra

| **Frame** | **ds*TcVER* median $\rho$** | **ds*TcFABP8* median $\rho$** | **ds*TcVER* mean $\rho$** | **ds*TcFABP8* mean $\rho$** |
|---|---|---|---|---|
| A | 0.121 | 0.099 | 0.173 | 0.113 |
| B | 0.185 | 0.116 | 0.187 | 0.118 |
| C | 0.110 | 0.062 | 0.160 | 0.072 |
